# Depletion of chemoresponsive mitochondrial fission mediator DRP1 does not mitigate sarcoma resistance

**DOI:** 10.1101/2024.06.05.597619

**Authors:** Karolina Borankova, Matyas Solny, Maria Krchniakova, Jan Skoda

**Author notes:** **Corresponding Author:** Dr. Jan Skoda, Department of Experimental Biology, Faculty of Science, Masaryk University University Campus Bohunice - C13.326, Kamenice 5, 625 00 Brno, Czech Republic. **Competing interests:** The authors declare no competing financial interests.

## Abstract

Specific patterns of mitochondrial dynamics have been repeatedly reported to promote drug resistance in cancer. However, whether targeting mitochondrial fission- and fusion-related proteins could be leveraged to combat multidrug-resistant pediatric sarcomas is poorly understood. Here, we demonstrated that the expression and activation of the mitochondrial fission mediator DRP1 are affected by chemotherapy exposure in common pediatric sarcomas, i.e., rhabdomyosarcoma and osteosarcoma. Unexpectedly, decreasing DRP1 activity through stable DRP1 knockdown did not attenuate sarcoma drug resistance or affect growth rate or the mitochondrial network morphology. The minimal impact on sarcoma cell physiology, combined with the upregulation of fission adaptor proteins (MFF and FIS1) detected in rhabdomyosarcoma cells, indicates that an alternative DRP1-independent mitochondrial fission mechanism may efficiently compensate for the lack of DRP1 activity. By exploring the upstream mitophagy and mitochondrial fission regulator, AMPKα1, we found that markedly reduced AMPKα1 levels are sufficient to maintain AMPK signaling capacity without affecting chemosensitivity. Collectively, our findings challenge the direct involvement of DRP1 in pediatric sarcoma drug resistance and highlight the complexity of yet-to-be-characterized noncanonical regulators of mitochondrial dynamics.

**SUMMARY BLURB:** The mitochondrial fission mediator DRP1 levels and activation are modulated upon chemotherapy exposure, yet depleting DRP1 does not restore chemosensitivity in the most common pediatric sarcomas.

## 1. INTRODUCTION

Pediatric sarcomas are a group of distinct mesenchymal solid tumors that arise either in bone or in mesenchymal soft tissue and account for approximately 20% of childhood nonhematologic malignancies (1). Therapy failure due to the development of multidrug resistance, resulting in a dismal prognosis, is frequent in the most prevalent pediatric sarcomas, such as rhabdomyosarcoma and osteosarcoma; the 5-year survival rates of their high-grade forms reach less than 50% (2,3,4). Although specific patterns of mitochondrial dynamics have emerged as important contributors to multidrug resistance in various cancers (5,6,7), little is known about the role of mitochondrial dynamics in pediatric sarcomas.

Mitochondrial dynamics is an umbrella term for mitochondrial adaptations consisting of mitochondrial fission and fusion, which shape the mitochondrial network, and mitochondrial biogenesis and mitophagy, a selective mitochondrial degradation process that fine-tunes mitochondrial quantity and quality (7). Deregulated mitochondrial dynamics have been implicated in tumorigenesis, as the expression of mitochondrial dynamics-related proteins differs between tumor and adjacent non-tumor tissues (8,9,10,11,12) and can be applied to predict patients’ outcomes (11,13,14,15,16,17). Furthermore, enhanced mitochondrial fission (18,19,20,21) and mitophagy (20,21,22,23) have been shown to promote a cancer stem-like phenotype in various tumor types. Consistently, mitochondrial fission (17,24,25,26,27) and fusion (28,29,30,31,32), as well as mitophagy (33,34,35), have been reported to mediate multidrug resistance, which can be mitigated by both pharmacological inhibition and modulation of the expression of mitochondrial dynamics-related proteins. Together, these reports suggest mitochondrial dynamics as a promising therapeutic target for overcoming resistance to standard treatment protocols. However, discrepancies between individual reports and contradictory results in various models indicate vast complexity and high context dependency, hindering translation into the clinic. For instance, shifting mitochondrial dynamics toward fission (24,28,29,30,31,32) or fusion (9,25,26,27,36) has been reported to suppress resistance to cancer therapy. Therefore, further research is needed to elucidate the tumor type-specific role of mitochondrial dynamics in multidrug resistance and its implications for cancer.

To investigate the potential of targeting mitochondrial dynamics for sarcoma therapy, we analyzed the modulation of mitochondrial fission and fusion-related proteins following drug exposure in the most common pediatric sarcomas, rhabdomyosarcoma and osteosarcoma, which have been largely overlooked in mitochondrial research thus far. Notably, we showed that the expression and activating phosphorylation of the mitochondrial fission mediator DRP1 are modulated in sarcoma cells upon chemotherapy exposure. However, depletion of DRP1 or the mitochondrial dynamics upstream regulator AMP-activated protein kinase catalytic subunit alpha 1 (AMPKα1) strikingly did not affect sarcoma cell physiology or attenuate multidrug resistance. These results provide new mechanistic evidence of noncanonical DRP1-independent mitochondrial fission, revealing the potential of sarcoma cells to escape DRP1-targeted anticancer therapies.

## 2. RESULTS

### 2.1. The mitochondrial fission mediator DRP1 predicts outcomes in transcriptomic cohorts and exhibits enhanced activation in post-therapy pediatric sarcoma-derived cells

To explore the therapeutic potential of mitochondrial dynamics-related proteins (**Fig. 1a**), we first analyzed the expression of their respective genes in relation to patient survival using publicly available rhabdomyosarcoma and osteosarcoma transcriptomics datasets. This analysis revealed that the expression of the gene encoding the mitochondrial fission mediator DRP1, *DNM1L,* could stratify both the rhabdomyosarcoma and osteosarcoma cohorts into groups with significant differences in survival (**Fig. 1b**). Interestingly, high *DNM1L* expression was associated with a poor prognosis in patients with osteosarcoma but predicted better outcomes in patients with rhabdomyosarcoma, suggesting a possible sarcoma subtype-dependent role for mitochondrial fission in the therapeutic response.

**Fig. 1:**
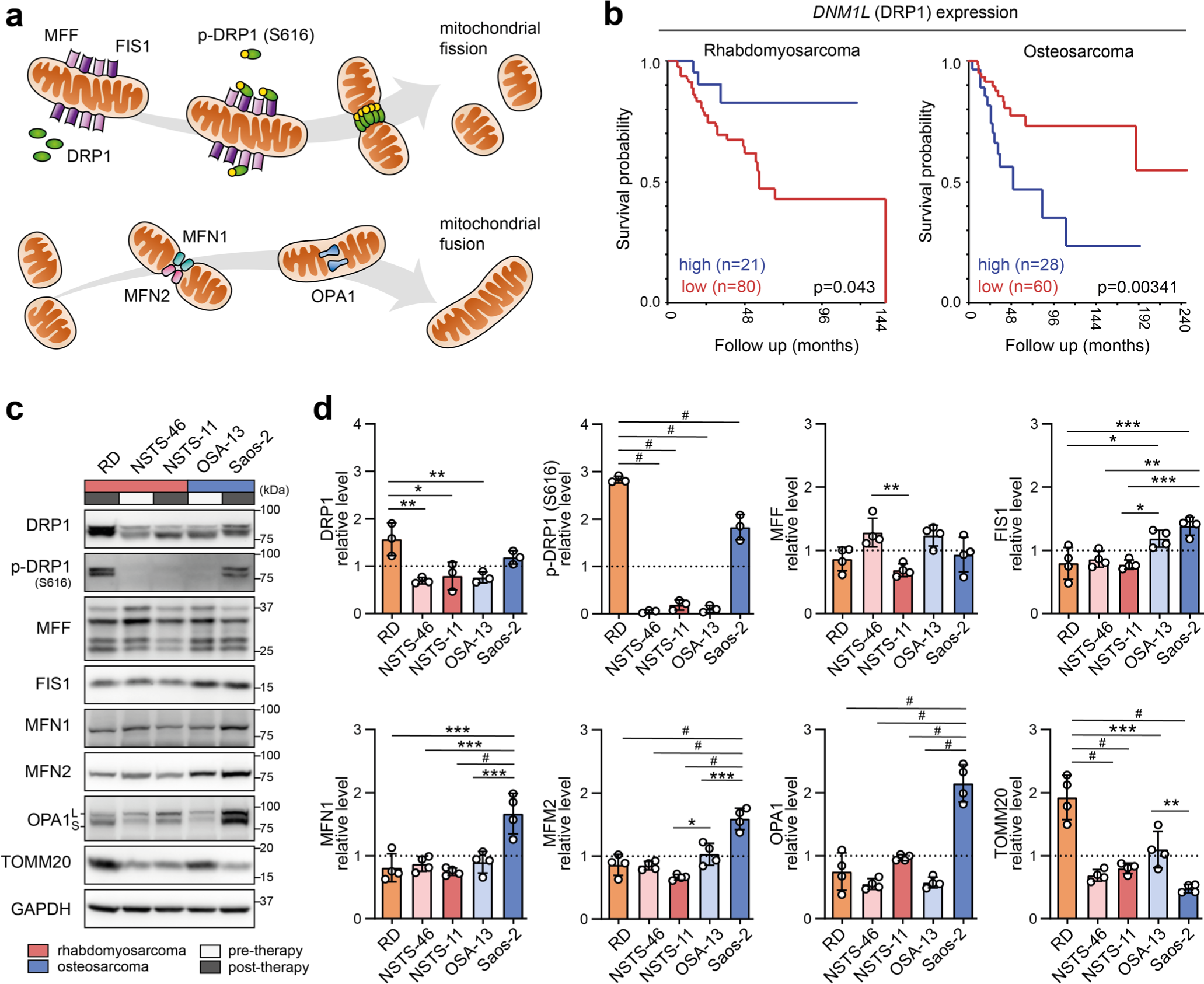
The mitochondrial fission mediator DRP1 predicts outcomes in transcriptomic cohorts and shows enhanced activation in post-therapy pediatric sarcoma-derived cells. (a) An overview of mitochondrial fission and fusion-related proteins. The mitochondrial fission mediator DRP1, which is activated by the phosphorylation of S616, forms a contractile ring on the mitochondrial surface by interacting with adaptor proteins such as mitochondrial fission factor (MFF) and FIS1, leading to mitochondrial constriction and fission. Outer mitochondrial membrane fusion is mediated by mitofusin-1 (MFN1) and mitofusin-2 (MFN2). Fusion of the inner mitochondrial membrane is mediated by OPA1. (b) Kaplan‒Meier analysis of the overall survival of patients stratified by the expression of the DRP1 gene, *DNML1*, was performed via the R2: Genomics Analysis and Visualization Platform. The analysis of rhabdomyosarcoma (76) and osteosarcoma (75) transcriptomic datasets was performed using scan cutoff mode with a minimal group size of 20 patients. (c, d) Western blotting (c) and densitometric analysis (d) of mitochondrial fission and fusion-related proteins across a panel of rhabdomyosarcoma and osteosarcoma cell lines derived from both therapy-naïve (pre-therapy) and relapsed/refractory (post-therapy) tumors. Statistical significance was determined by one-way ANOVA followed by Tukey’s multiple comparisons test (d), *p<0.05, **p<0.01, ***p<0.001, #p<0.0001.

To elucidate whether initial drug exposure affects mitochondrial dynamics, we included rhabdomyosarcoma and osteosarcoma cell lines derived from both therapy-naïve (pre-therapy) and relapsed/refractory (post-therapy) tumors. Immunoblotting revealed differential expression of mitochondrial fission- and fusion-related proteins across a panel of tested sarcoma cells (**Fig. 1c, d**). Notably, the activating phosphorylation of the mitochondrial fission mediator DRP1 on serine 616 (37) was strongly upregulated in 2 out of 3 post-therapy cell lines, indicating that enhanced DRP1 activation is a potential adaptation to therapy-induced stress. Interestingly, post-therapy osteosarcoma Saos-2 cells showed upregulated levels of both mitochondrial fission- and fusion-related proteins despite maintaining relatively low mitochondrial mass, as assessed by the expression of the mitochondrial import receptor subunit TOMM20, which serves as a proxy for mitochondrial content. In contrast, the post-therapy RD rhabdomyosarcoma cell line exhibited increased levels solely in the mitochondrial fission mediator DRP1, along with strong upregulation of its activating phosphorylation. Given these potential therapy-related patterns, we aimed to analyze whether these differences are reflected in the modulation of mitochondrial fission and fusion-related proteins in response to chemotherapy exposure.

### 2.2. Autophagy inhibition suppresses the drug-induced activating phosphorylation of DRP1 but does not mitigate rhabdomyosarcoma drug resistance

Mitochondrial fission mediated by DRP1 not only changes mitochondrial morphology but also separates damaged mitochondria into autophagosomes for autophagic degradation (38). The ongoing autophagic flux can be pharmacologically blocked by bafilomycin A1 (**Fig. 2a**). Compared with other sarcoma-derived cell lines, rhabdomyosarcoma RD cells, which exhibit high levels of DRP1 expression and its activating phosphorylation, exhibited greater vulnerability to bafilomycin A1-mediated inhibition of autophagy (**Fig. 2b**). Considering the high sensitivity to autophagy inhibition and the increased levels of DRP1 activating phosphorylation, we assumed that high autophagic flux, which indicates high mitophagy activity, is crucial for the survival of RD cells. Hence, mitochondrial fission and fusion-related protein levels were analyzed in RD cells after exposure to drugs at half-maximal inhibitory concentrations (IC_50_ values, dose‒response curves detailed in **Fig. S1**) and autophagy inhibition by a sublethal dose of bafilomycin A1. Standard chemotherapy drugs commonly used for pediatric sarcoma—cisplatin (39,40), doxorubicin (40,41), and topotecan (42,43), all of which have predominantly genotoxic effects, and a microtubule polymerization inhibitor, vincristine (41,44)—were used in these experiments. Exposure to chemotherapeutics *in vitro* increased the level of DRP1 activating phosphorylation, and this effect was diminished by concomitant autophagy inhibition with bafilomycin A1 (**Fig. 2c, d**). This finding suggested that the activation of DRP1-mediated mitochondrial fission serves as an adaptation to chemotherapy-induced stress and that inhibiting autophagy, thereby mitophagy, prevents this response.

**Fig. 2:**
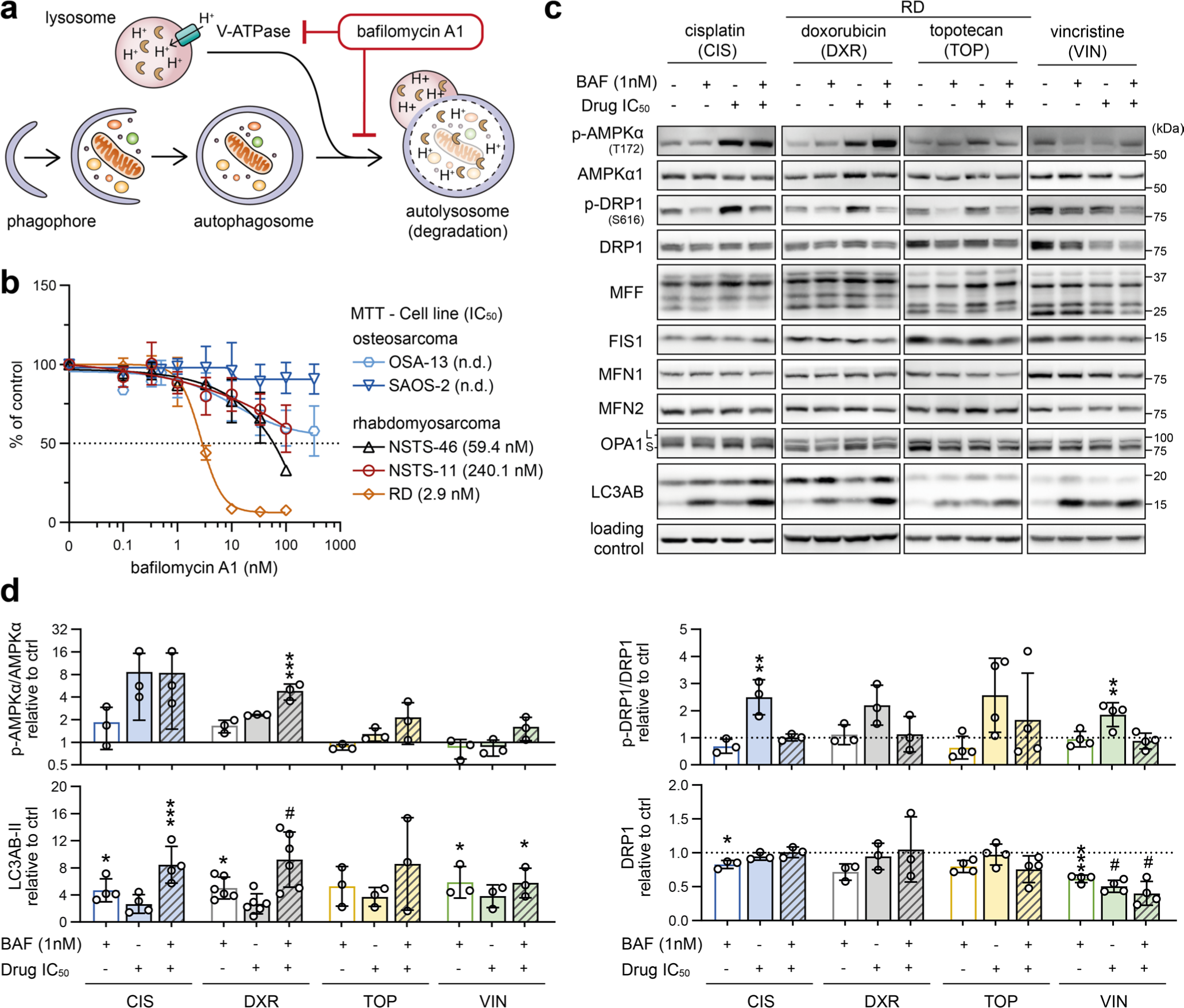
Bafilomycin A1-mediated autophagy inhibition suppresses the drug-induced activating phosphorylation of DRP1. **(a)** An overview of the mechanism of bafilomycin A1-mediated autophagy inhibition. Cellular components marked for autophagic degradation are engulfed by the phagophore, leading to autophagosome formation. The autophagosome content is degraded due to fusion with acidified lysosomes. Bafilomycin A1 inhibits both lysosomal acidification and lysosome-autophagosome fusion, thereby inhibiting autophagic flux. **(b)** MTT viability assay analysis after 72 hours of treatment revealed variable sensitivity to bafilomycin A1 in a panel of rhabdomyosarcoma and osteosarcoma cell lines derived from both therapy-naïve (pre-therapy) and relapsed/refractory (post-therapy) tumors. The respective IC50 values are indicated in brackets; n.d. indicates that the IC50 could not be determined due to marked resistance. The data points represent the mean ± SD, biological n=3, technical n=3. **(c, d)** Western blotting (c) and densitometric analysis (d) of proteins related to mitochondrial fission/fusion and autophagy in RD rhabdomyosarcoma cells exposed to 1 nM bafilomycin A1 (BAF), chemotherapy drugs at the IC50 doses or combinations of BAF with chemotherapy drugs for 72 hours. Normalized protein levels are plotted relative to the untreated controls; mean ± SD. BAF – bafilomycin A1, CIS – cisplatin, DXR – doxorubicin, TOP – topotecan, VIN – vincristine. Densitometric analysis of MFF, FIS1, MFN1, MFN2, and OPA1 is provided in **Fig. S2**. Statistical significance was determined by one-way ANOVA followed by Tukey’s multiple comparisons test (d), *p<0.05, **p<0.01, ***p<0.001, #p<0.0001.

Unexpectedly, other mitochondrial fission- and fusion-related proteins in RD cells were generally unaffected by drug exposure (**Figs. 2c, S2**). However, the stress-responsive AMP-activated protein kinase (AMPK), known to promote both autophagy and mitochondrial fission (45), was activated by cisplatin and doxorubicin treatment (**Fig. 2c, d**). Considering that both cisplatin and doxorubicin enhanced autophagic flux in RD cells, as indicated by the increase in the LC3AB-II isoform upon bafilomycin A1-mediated inhibition of autophagy (**Fig. 2c, d**), we speculated that, in line with reports in other cell types (33,46,47), autophagy might protect rhabdomyosarcoma cells against these genotoxic drugs. To test this hypothesis, we treated rhabdomyosarcoma cells with cisplatin and doxorubicin as single agents or in combination with the autophagy inhibitor bafilomycin A1 or the autophagy inducer rapamycin. Rapamycin is known to promote autophagy by inhibiting mammalian target of rapamycin complex 1 (mTORC1), a central controller of cell metabolism and a key negative regulator of the ULK1 autophagy initiation complex (48). Indeed, rapamycin reduced sensitivity to cisplatin and doxorubicin in both bafilomycin A1-sensitive (RD) and bafilomycin A1-resistant (NSTS-46 and NSTS-11) rhabdomyosarcoma cells (**Fig. 3**), suggesting that enhanced autophagy might serve as a mechanism of multidrug resistance in rhabdomyosarcoma. However, combined treatment with bafilomycin A1 did not enhance the effects of these chemotherapy drugs, even in autophagy inhibition-sensitive RD cells (**Fig. S3**). This finding indicates that the autophagy-independent effects of mTORC1 inhibition promote the resistance of rhabdomyosarcoma cells to cisplatin and doxorubicin. These results additionally contradict the cytoprotective role of drug-induced activation of DRP1, which was diminished by bafilomycin A1.

**Fig. 3:**
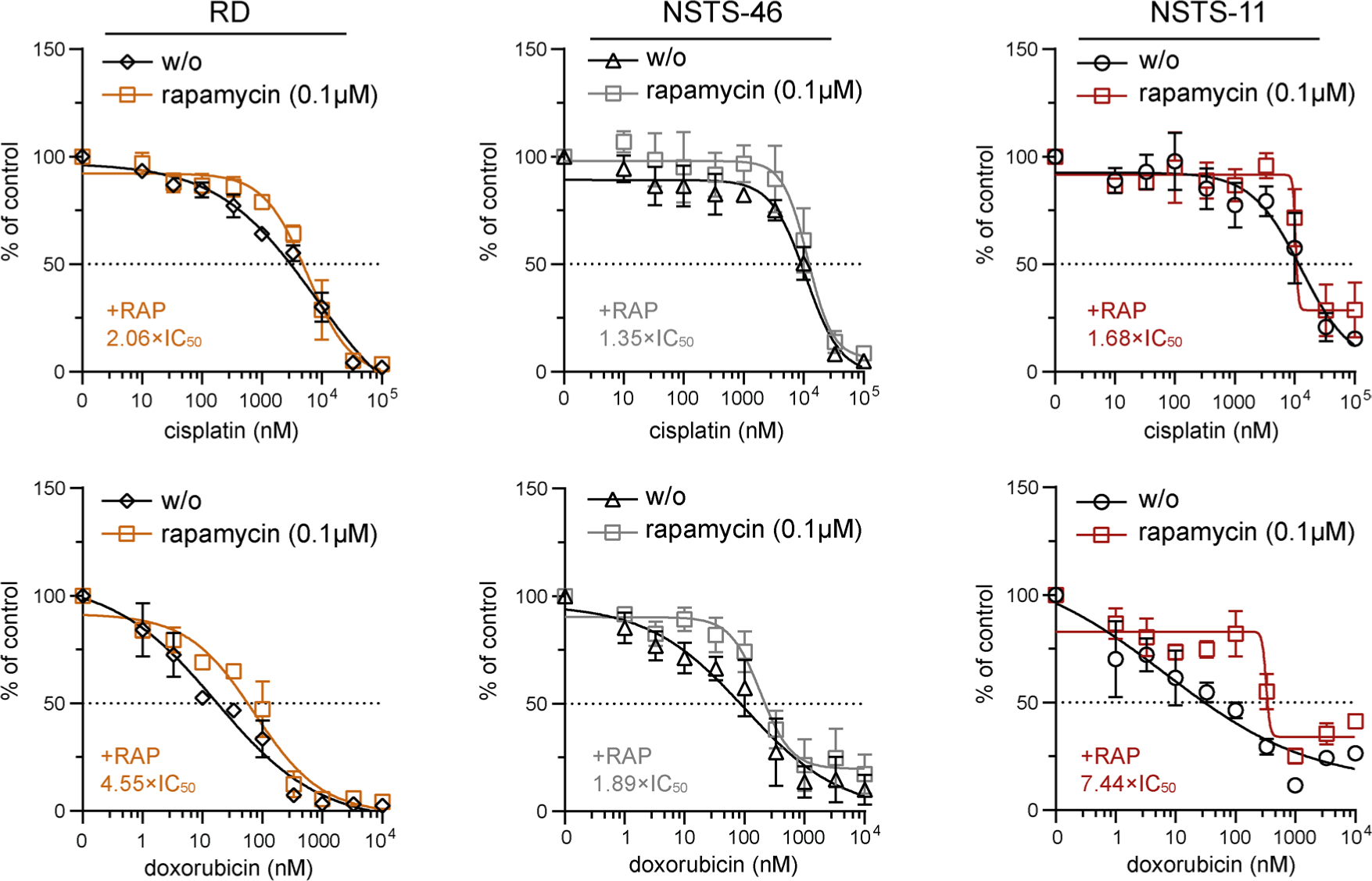
The mTOR inhibitor rapamycin reduced sensitivity to cisplatin and doxorubicin in both bafilomycin A1-sensitive and bafilomycin A1-resistant rhabdomyosarcoma cells. MTT viability assay analysis of bafilomycin A1-sensitive (RD) and bafilomycin A1-resistant (NSTS-46, NSTS-11) rhabdomyosarcoma cells exposed to the indicated concentrations of cisplatin and doxorubicin alone or in the presence of 0.1 μM rapamycin for 72 hours. The rapamycin-induced increase in the IC50 is indicated as the ratio of the IC50 determined for the combination of rapamycin and chemotherapy drugs to the IC50 determined for chemotherapy drugs alone. The data points represent the mean ± SD, biological n≥3, technical n=3.

### 2.3. Modulation of the mitochondrial fission machinery in sarcoma cells upon chemotherapy exposure reflects the initial DRP1 status

Analysis of mitochondrial fission and fusion-related proteins in bafilomycin-resistant rhabdomyosarcoma cells (**Figs. 4a, c, S4**) and osteosarcoma cells (**Figs. 4b, c, S4**) exposed to IC_50_ doses of chemotherapy drugs (as detailed in **Fig. S1**) revealed that the drug-induced modulation of DRP1 depended on the initial status of DRP1. In Saos-2 osteosarcoma cells with increased levels of DRP1 activating phosphorylation (**Fig. 1c, d**), exposure to genotoxic drugs increased the levels of this p-DRP1(S616) form (**Fig. 4b, c**). In contrast, in sarcoma cells without detectable p-DRP1(S616), chemotherapy exposure further suppressed the levels of mitochondrial fission machinery proteins (**Fig. 4a, c**). A decrease in total DRP1 was detected in NSTS-11 cells, whereas NSTS-46 and OSA-13 cells, which expressed the highest basal levels of the DRP1 adaptor MFF (**Fig. 1c, d**), showed prevalent downregulation of MFF in response to the tested drugs (**Fig. 4a-c**). Interestingly, the levels of FIS1, another DRP1 adaptor protein, did not show any clear trend upon drug exposure (**Fig. S4**), suggesting that only a subset of mitochondrial fission-related proteins are affected by drug treatment. Conversely, the expression of fusion-related proteins was mostly unaffected after treatment (**Fig. 4a-c, S4**), with one exception: mitofusin-1 (MFN1) was downregulated in response to topotecan in NSTS-11 rhabdomyosarcoma cells and both osteosarcoma cell lines. While these results exclude the role of MFN1 in sarcoma multidrug resistance, further research might elucidate its potential involvement in regulating selective resistance to topotecan. In addition to mitofusin-2 (MFN2), the mediator of inner mitochondrial membrane fusion, OPA1, remained intact in drug-exposed sarcoma cells (**Fig. 4a, b, S4**), demonstrating that chemotherapy did not activate the mitochondrial metalloproteases OMA1 and YME1L1, which cleave OPA1 into its short (S-OPA1) forms under mitochondrial stress. Unexpectedly, we observed preferential downregulation of these mitochondrial stress sensors in drug-exposed sarcoma cells (**Fig. S5**), suggesting that OMA1 and YME1L1 may have OPA1 processing-independent functions in sarcoma chemotherapy adaptation; similarly, OMA1 was recently reported to induce a metabolic shift upon DNA damage (49).

**Fig. 4:**
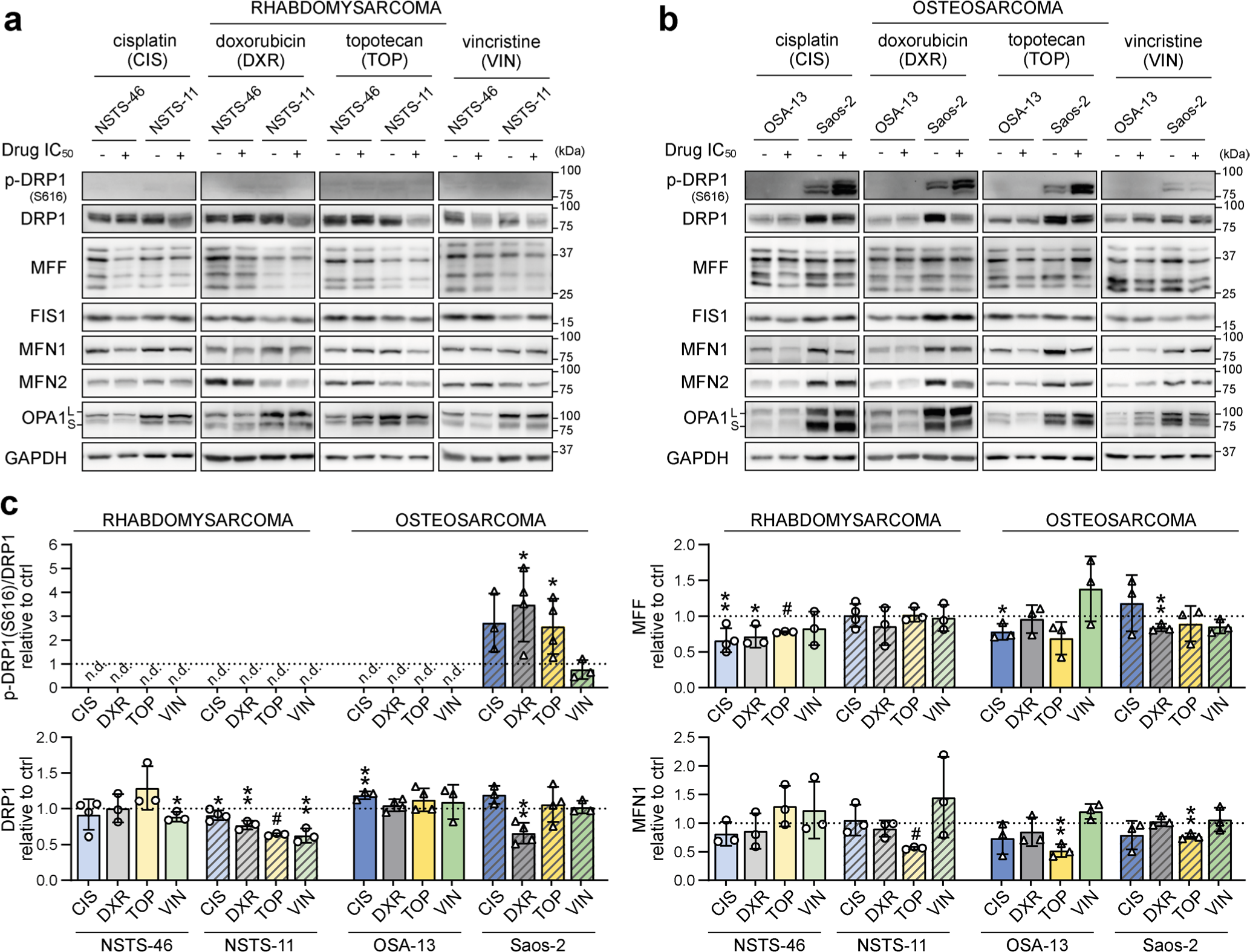
Modulation of the mitochondrial fission machinery upon chemotherapy exposure reflects the initial DRP1 status in sarcoma cells. **(a-c)** Western blotting (a, b) and densitometric analysis (c) of mitochondrial fission and fusion-related proteins in rhabdomyosarcoma (a, c) and osteosarcoma (b, c) cells exposed to IC50 doses of chemotherapy for 72 hours. Normalized protein levels are plotted relative to the untreated controls as the mean ± SD. CIS – cisplatin, DXR – doxorubicin, TOP – topotecan, VIN – vincristine. Densitometric analysis of FIS1, MFN2, and OPA1 is provided in **Fig. S4**. Statistical significance was determined by unpaired two-tailed Student’s t-test (c), *p<0.05, **p<0.01, #p<0.0001, n.d. – not detected.

### 2.4. Stable knockdown of DRP1 does not impair sarcoma cell physiology or decrease drug resistance

Collectively, the analysis of mitochondrial dynamics following drug exposure suggested that mitochondrial fission is a context-dependent mechanism that fine-tunes drug resistance. Since treatment with chemotherapeutics increased DRP1 activating phosphorylation in post-therapy sarcoma cells, we hypothesized that DRP1 activity protects these cells against chemotherapy-induced stress. To test this hypothesis, we established cells with stable DRP1 knockdown by lentiviral transduction of constructs encoding target-specific shRNAs under the control of a constitutive promoter. Characterization of single-cell-derived clones from DRP1-knockdown (shDRP1) and control (shCTRL) cells confirmed the depletion of DRP1 and demonstrated a marked decrease in its activating phosphorylation form in both rhabdomyosarcoma and osteosarcoma models (**Figs. 5a, e, S6**). Despite significantly decreased DRP1 activity, the growth rates of rhabdomyosarcoma (**Fig. 5b**) and osteosarcoma (**Fig. 5f**) shDRP1 clones were very similar to those of their control counterparts, suggesting that DRP1 is not required for sarcoma cell cycle progression. Strikingly, DRP1 downregulation did not mitigate chemoresistance in either rhabdomyosarcoma or osteosarcoma models. A comparison of IC_50_ values determined in shCTRL and shDRP1 clones did not reveal any significant differences in resistance to standard chemotherapy (**Figs. 5c, g, S7a, S8a**) or targeted inhibitors (**Figs. 5d, h, S7b, S8b**), except for slightly reduced sensitivity to cisplatin and vincristine detected in rhabdomyosarcoma but not osteosarcoma shDRP1 clones (**Figs. 5c, g, S7, S8**). Hence, we hypothesized that DRP1 plays a dispensable role in sarcoma cell physiology, which is further supported by the similar sensitivities of the shCTRL and shDRP1 clones to autophagy inhibitors (bafilomycin A1 and chloroquine) and the respiratory complex I inhibitor phenformin (**Fig. S9**).

**Fig. 5:**
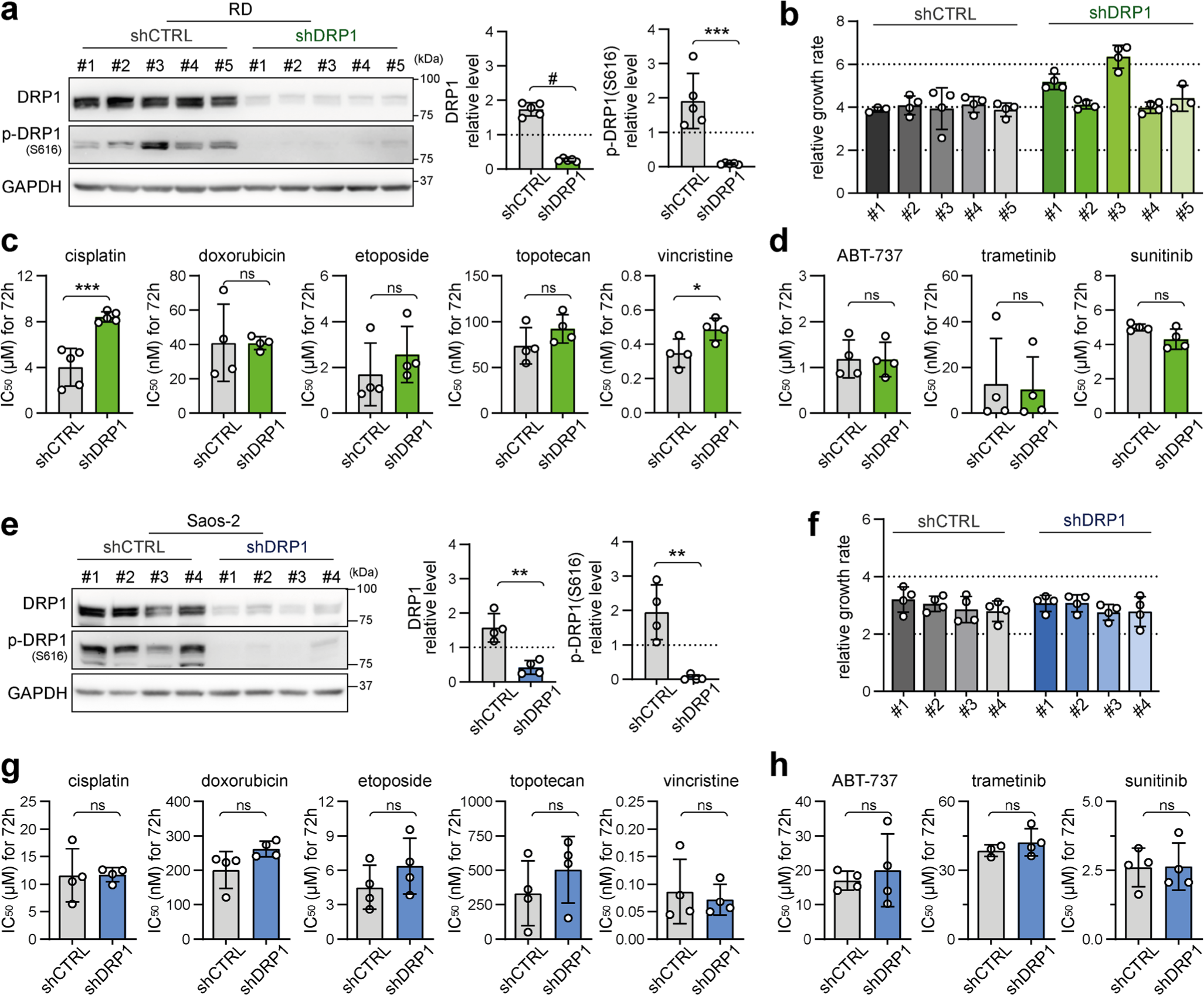
Stable knockdown of DRP1 does not impair sarcoma cell growth or enhance drug sensitivity. **(a, e)** Western blotting and densitometric analysis confirmed the efficiency of DRP1 knockdown in rhabdomyosarcoma RD (a) and osteosarcoma Saos-2 (e) cell clones. The mean normalized protein levels detected in the control (shCTRL) and DRP1-downregulated (shDRP1) cell clone groups are plotted as the mean ± SD. Densitometric analysis of biological replicates of individual cell clones is provided in **Fig. S6a**. **(b, f)** Live-cell imaging analysis of the growth rate of individual RD (b) and Saos-2 (f) shCTRL and shDRP1 cell clones. The data are plotted as the mean ± SD of the fold change in cell confluence after 72 h of incubation. **(c, d, g, h)** The chemoresistance of control and DRP1-downregulated RD cells was compared using the drug IC50. The data are plotted as the mean ± SD of the IC50 values determined for individual RD (c, d) and Saos-2 (g, h) shCTRL and shDRP1 cell clones. Dose‒response curves for standard chemotherapy drugs (c, g) and targeted inhibitors (d, h) are provided in **Fig. S7; S8.** Statistical significance was determined by unpaired two-tailed Student’s t-test, *p<0.05, **p<0.01, ***p<0.001, #p<0.0001, ns – not significant.

**Fig. 6:**
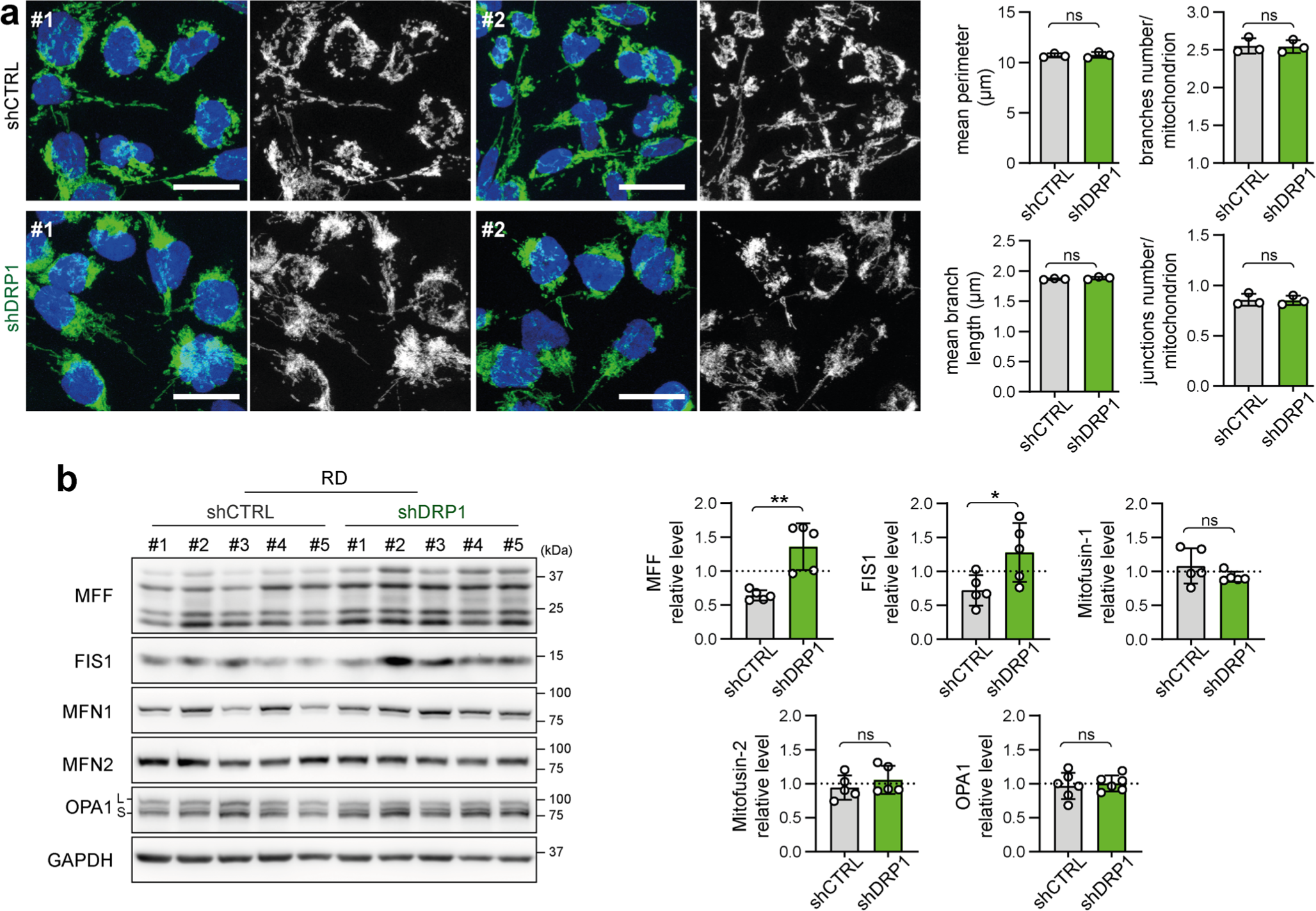
Depletion of DRP1 does not affect the mitochondrial network morphology in RD rhabdomyosarcoma cells. **(a)** Mitochondrial morphology, as visualized by immunofluorescence staining of TOMM20 (green), was not altered by DRP1 downregulation. Representative merged images (green – TOMM20, blue – nuclei) and grayscale images (mitochondria) of maximum intensity projections of confocal microscopy Z-stacks are shown for individual RD control (shCTRL) and DRP1-downregulated (shDRP1) cell clones. White bar – 20 μm. Image analysis using the ImageJ plug-in tool Mitochondria Analyzer showed that parameters describing the mitochondrial network morphology did not significantly differ between the shCTRL and shDRP1 clones. The mean parameters determined for the shCTRL and shDRP1 clone groups are presented as the mean ± SD. Analysis of biological replicates of individual cell clones is provided in **Fig. S10**. **(b)** Western blotting and densitometric analysis of mitochondrial fission and fusion-related proteins in RD shCTRL and shDRP1 cell clones. The mean normalized protein levels detected in the shCTRL and shDRP1 clone groups are plotted relative to the average levels as the mean ± SD. Densitometric analysis of biological replicates of individual cell clones is provided in **Fig. S12**. Statistical significance was determined by unpaired two-tailed Student’s t-test, *p<0.05, **p<0.01, ns – not significant.

**Fig. 7:**
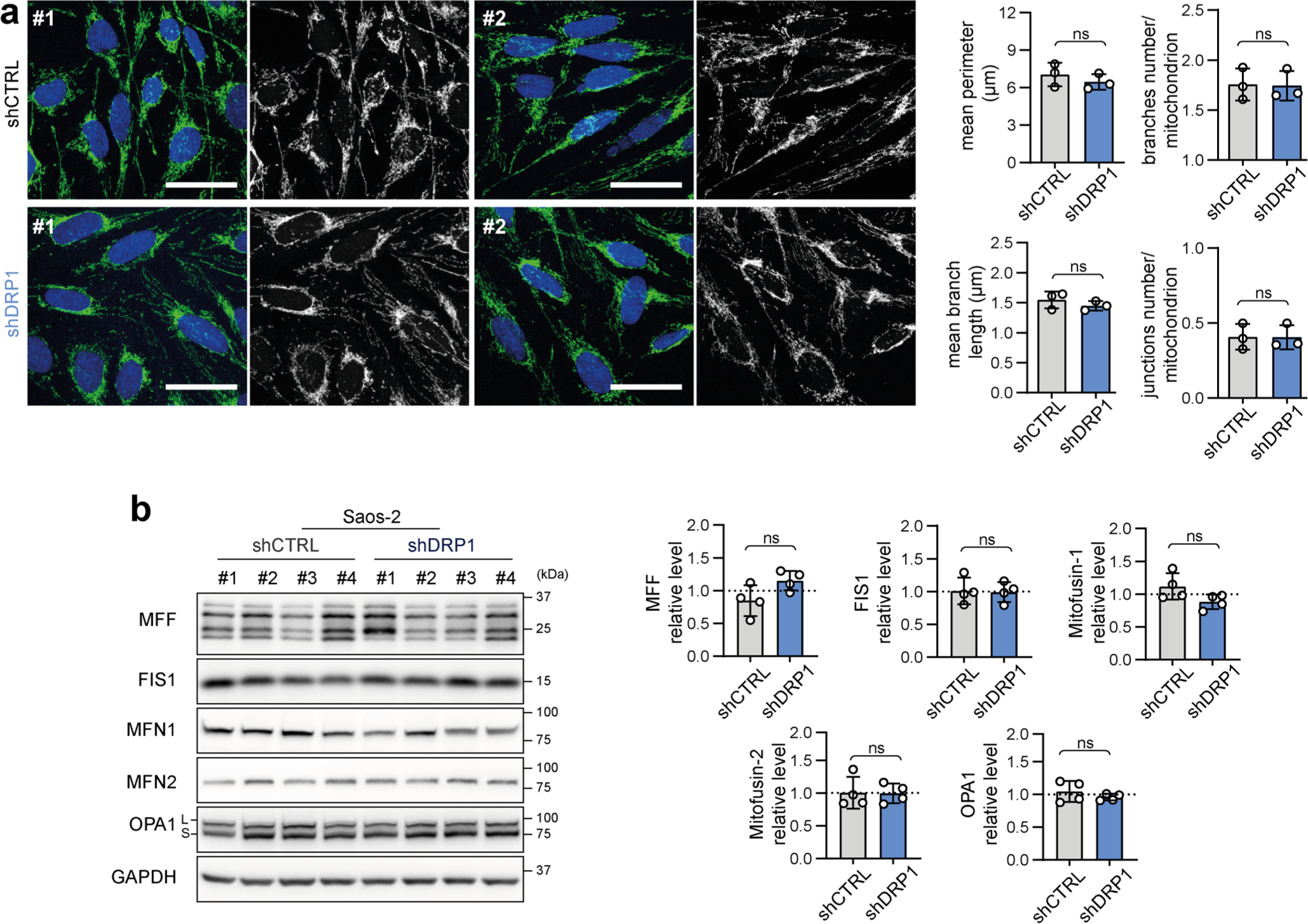
Depletion of DRP1 does not affect the mitochondrial network morphology in Saos-2 osteosarcoma cells. **(a)** Mitochondrial morphology, as visualized by immunofluorescence staining of TOMM20 (green), was not altered by DRP1 downregulation. Representative merged images (green – TOMM20, blue – nuclei) and grayscale images (mitochondria) of maximum intensity projections of confocal microscopy Z-stacks are shown for individual Saos-2 control (shCTRL) and DRP1-downregulated (shDRP1) cell clones. White bar – 40 μm. Image analysis using the ImageJ plug-in tool Mitochondria Analyzer showed that parameters describing the mitochondrial network morphology did not significantly differ between the shCTRL and shDRP1 clones. The mean parameters determined for the shCTRL and shDRP1 clone groups are presented as the mean ± SD. Analysis of biological replicates of individual cell clones is provided in **Fig. S11**. **(b)** Western blotting and densitometric analysis of mitochondrial fission and fusion-related proteins in Saos-2 shCTRL and shDRP1 cell clones. Normalized protein levels detected in the shCTRL and shDRP1 clone groups are plotted relative to the average level as the mean ± SD. Densitometric analysis of biological replicates of individual cell clones is provided in Fig. **S13**. Statistical significance was determined by unpaired two-tailed Student’s t-test, ns – not significant.

**Fig. 8:**
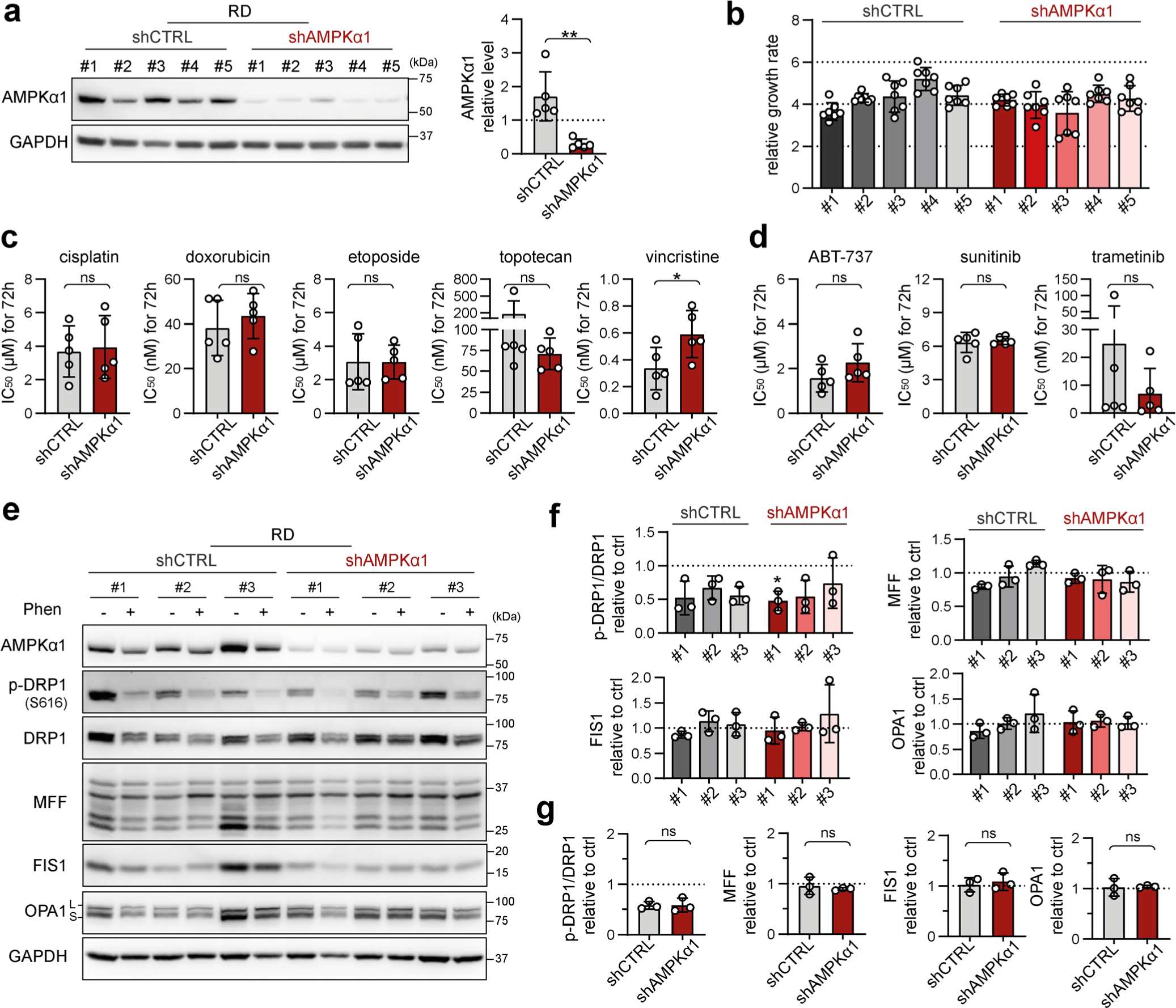
Rhabdomyosarcoma drug resistance and stress-induced modulation of mitochondrial dynamics-related proteins are independent of the upstream mitochondrial fission and autophagy inducer AMPKα1. **(a)** Western blotting and densitometric analysis confirmed the efficiency of AMPKα1 knockdown in rhabdomyosarcoma RD cell clones. The mean normalized protein levels detected in the control (shCTRL) and AMPKα1-downregulated (shAMPKα1) cell clone groups are plotted relative to the average level as the mean ± SD. Densitometric analysis of biological replicates of individual cell clones is provided in **Fig. S14**. **(b)** Live-cell imaging analysis of the growth rate of individual shCTRL and shAMPKα1 RD cell clones. The data are plotted as the mean ± SD of the fold change in cell confluence after 72 hours of incubation. **(c, d)** The chemoresistance of control and AMPKα1-downregulated RD cells was compared using the drug IC50. The data are plotted as the mean ± SD of the IC50 values determined for individual shCTRL and shAMPKα1 cell clones. Dose‒response curves for standard chemotherapy drugs (c) and targeted inhibitors (d) are provided in **Fig. S15**. **(e-g)** Western blotting (e) and densitometric analysis (f, g) of mitochondrial fission- and fusion-related proteins in rhabdomyosarcoma RD shCTRL and shAMPKα1 cell clones exposed for 72 hours to 1 mM phenformin. Normalized protein levels detected in individual cell clones are plotted relative to the untreated controls as the mean ± SD (f). Comparison of the phenformin-induced modulation of mitochondrial fission and fusion-related proteins detected in the RD shCTRL and shAMPKα1 cell clone groups (g). Average relative protein levels per group and biological replicate are plotted as the mean ± SD. Densitometric analysis of biological replicates of individual cell clones and extended analysis of fusion-related proteins after 72 hours of phenformin treatment are provided in **Fig. S17**. Statistical significance was determined by unpaired two-tailed Student’s t-test (c, d, g) or by one-way ANOVA followed by Tukey’s multiple comparisons test (f), *p<0.05, ns – not significant.

To further elucidate the very limited impact of DRP1 knockdown on sarcoma cell physiology, we aimed to assess whether DRP1 depletion shifted the balance between mitochondrial fission and fusion. Microscopically, the morphology of the mitochondrial network was unchanged in the shDRP1 clones compared to the control clones in both rhabdomyosarcoma (**Figs. 6a, S10**) and osteosarcoma (**Figs. 7a, S11**). Considering the near absence of the activated p-DRP1(S616) form in shDRP1 clones, these results indicate that DRP1-independent mitochondrial fission compensates for the lack of mitochondrial membrane scission mediated by DRP1. In the rhabdomyosarcoma model, immunoblotting revealed upregulation of the mitochondrial outer membrane adaptor proteins MFF and FIS1 (**Figs. 6b, S12**), which might contribute to alternative DRP1-independent mitochondrial fission (50). Nevertheless, further research is needed to elucidate the mechanisms compensating for DRP1 inactivation because no modulation of mitochondrial fission or fusion-related proteins was detected in osteosarcoma shDRP1 clones (**Figs. 7b, S13**).

### 2.5. Rhabdomyosarcoma drug resistance and stress-induced modulation of mitochondrial dynamics proteins are independent of the upstream mitochondrial fission and autophagy inducer AMPKα1

Given the identified capacity of sarcoma cells to compensate for DRP1, which is presumably a key component of the mitochondrial fission machinery, we hypothesized that targeting upstream regulators might be necessary to overcome drug resistance. Therefore, we established rhabdomyosarcoma clones with stable knockdown of AMPKα1 (**Figs. 8a, S14**), an isoform of the catalytic subunit of stress-responsive AMPK that promotes both autophagy and mitochondrial fission upon activation (45). This AMPKα1 isoform, which is frequently amplified in tumors (51,52), has been reported to enhance resistance against genotoxic agents (53,54). Notably, we demonstrated that cisplatin and doxorubicin treatments activated AMPK in RD cells (**Fig. 2c, d**). As expected, no difference in growth rate was detected between the control (shCTRL) and AMPKα1-knockdown (shAMPKα1) clones (**Fig. 8b**), as AMPK activity is induced solely under stress conditions. However, the downregulation of AMPKα1 did not decrease resistance to standard chemotherapy (**Figs. 8c, S15**) or targeted inhibitors (**Figs. 8d, S15**). Consistent with these findings, sensitivity to autophagy inhibitors (**Fig. S16**) and the respiratory complex I inhibitor phenformin (**Fig. S16**) was not affected by AMPKα1 knockdown.

Since drug resistance was unaffected by AMPKα1 knockdown, we aimed to investigate whether the downregulation of AMPKα1 modified mitochondrial dynamics downstream of activated AMPK. Unexpectedly, after 72 hours of exposure to a well-established AMPK activator, phenformin, no significant differences in the levels of mitochondrial fission or fusion-related proteins were detected between the shCTRL and shAMPKα1 clones (**Figs. 8e-g, S17**). In addition, upon nutrient deprivation combined with phenformin treatment, AMPKα1-knockdown cells exhibited a rapid increase in the level of activated AMPK, p-AMPKα(T172), which was comparable to that of their control counterparts (**Fig. S18**). Another AMPKα isoform, AMPKα2, might compensate for AMPKα1 depletion, potentially explaining the observed AMPK activation. However, this effect was excluded by immunoblotting, which demonstrated that AMPKα2 expression was undetectable in both the shCTRL and shAMPKα1 clones (**Fig. S14**), even upon phenformin-mediated AMPK activation (**Figs. S17, S18**). Collectively, our data reveal that very low levels of AMPK are sufficient to maintain AMPK signaling and mitochondrial responses after AMPKα1 knockdown. Thus, models with concurrent knockout of both AMPKα isoforms should provide a definite answer as to whether AMPK signaling might promote rhabdomyosarcoma multidrug resistance by regulating mitochondrial dynamics.

## 3. DISCUSSION

Drug resistance is a significant factor contributing to the failure of cancer therapy. However, effective treatment protocols that would prevent the induction of acquired resistance remain elusive. Recently, enhanced mitochondrial dynamics have emerged as a common vulnerability of drug-resistant and aggressive cancer cells, and the mitochondrial fission mediator DRP1 has been repeatedly suggested as a potential therapeutic target (9,11,17,18,27). High expression of DRP1 and its adaptors has been associated with poor survival in several adult tumor types, including breast carcinoma (13,55), nasopharyngeal carcinoma (17), lung carcinoma (11), and hepatocellular carcinoma (14,15,56,57). Consistent with these findings, we found that DRP1 expression can be used to stratify pediatric sarcoma patients according to survival, albeit in a sarcoma subtype-dependent manner (**Fig. 1b**). High DRP1 expression is associated with poor survival in patients with osteosarcoma but predicts better outcomes in patients with rhabdomyosarcoma, suggesting that mitochondrial fission plays a tumor type-dependent role in the therapeutic response. This finding in rhabdomyosarcoma aligns with studies that revealed an association between low expression of DRP1 adaptors and a poor prognosis in tongue squamous cell carcinoma patients (24,58). Moreover, similar findings have been observed in other malignancies, such as melanoma (59), lung adenocarcinoma (12), and hepatocellular carcinoma (10), where high expression of mitochondrial fusion-related genes is linked to a poor prognosis.

Although recent studies have demonstrated variable effects of chemotherapy on DRP1 (28,32,60) and other proteins involved in mitochondrial fission (24,58) and fusion (32,60,61), the underlying determinants are poorly understood. Using a panel of pediatric sarcoma cell lines with differential DRP1 expression and activation, we demonstrated that the chemotherapy-induced modulation of mitochondrial fission depends on the initial status of DRP1 (**Figs. 1c, d, 2c, d, 4a-c**). The activity of ERK was previously reported to enhance DRP1 activating phosphorylation (8,62). However, in RD cells, the increase in ERK activation did not correspond to the chemotherapy-induced activating phosphorylation of DRP1 (**Figs. 2c, d, S19**). Thus, we hypothesize that other putative mechanisms, such as DUSP6-mediated dephosphorylation of DRP1 at S616, may also contribute to the regulation of DRP1 phosphorylation in rhabdomyosarcoma (63).

Although depletion of DRP1 (27,55,56) or its adaptor protein MFF (25,26) has been shown to suppress drug resistance in carcinomas, stable DRP1 knockdown did not mitigate drug resistance in our sarcoma models (**Figs. 5, S7, S8**). Strikingly, DRP1 depletion even slightly increased resistance to cisplatin and vincristine in RD cells, likely indicating a predominant proapoptotic role of DRP1 in rhabdomyosarcoma, as found in other cancers (64,65). Notably, we observed that DRP1 knockdown had a very limited impact on sarcoma cell physiology. In contrast to reports indicating that DRP1 activity is necessary for proper cell cycle progression (66,67) and tumor growth (8,18,55), the growth rate of sarcoma cells was unaffected by reduced DRP1 levels (**Fig. 5b, f**). While previous studies suggested that DRP1 inhibits the cancer stem-like phenotype (18,68), we did not observe any downregulation of stemness-associated markers in shDRP1 rhabdomyosarcoma RD cells (**Fig. S20**). Importantly, DRP1 depletion in sarcoma models did not alter the mitochondrial network morphology (**Figs. 6a, 7a**), whereas numerous reports in other cell types have shown that DRP1 knockdown induces a more fused and less fragmented mitochondrial network (27,55,57,66,69). Taken together, our results suggest the presence of a potential DRP1-independent mechanism that may efficiently compensate for the lack of DRP1 activity in sarcoma cells. Although DRP1 activity is essential for canonical mitochondrial fission (70,71), dynamin-2 has been reported to be an alternative mediator of mitochondrial fission (72). Additionally, the DRP1 adaptor protein FIS1 has been shown to regulate mitochondrial fission by inhibiting mitochondrial fusion (50). Similar compensatory mechanisms might also be adopted by sarcoma cells, as evidenced by the upregulation of the DRP1 adaptor proteins MFF and FIS1 in DRP1-depleted rhabdomyosarcoma RD cells (**Figs. 6b, S12**), further suggesting that therapies targeting DRP1 for sarcoma may be likely to fail.

Overall, our study revealed that drug exposure modulates mitochondrial fission-related proteins in pediatric sarcomas. Importantly, we revealed a novel finding: the nature of chemotherapy-induced modulation appears to be influenced by the initial status of the DRP1 fission machinery. However, further research is needed to elucidate the possible clinical implications. Strikingly, stable DRP1 knockdown neither mitigated sarcoma chemoresistance nor impacted cell physiology, possibly due to the selection of cells capable of activating compensatory mechanisms. Thus, our results prompt investigations into noncanonical, DRP1-independent mitochondrial fission mechanisms, including upstream regulators, to better understand whether targeting mitochondrial dynamics could serve as a therapeutic strategy to overcome multidrug resistance.

## 4. MATERIALS AND METHODS

### 4.1. Cell culture and treatment

The following sarcoma cell lines were utilized in the study: (i) the osteosarcoma cell line OSA-13 and the embryonal rhabdomyosarcoma cell lines NSTS-11 and NSTS-46, all of which were derived in-house from tumor tissue with written informed consent under the IGA MZCR NR/9125-4 project approved by the Research Ethics Committee of the School of Medicine, Masaryk University, Brno, Czech Republic (approval no. 23/2005); and (ii) the osteosarcoma cell line Saos-2 and the rhabdomyosarcoma cell line RD, both of which were purchased from ECACC. In addition, the human embryonic kidney cell line HEK293T served as a positive control for detecting AMPKα2, as shown in **Figs. S14; S17; S18**. The authenticity of the cell lines was verified by STR profiling (Generi Biotech, Hradec Králové, Czech Republic), and all cell lines were regularly tested for mycoplasma contamination by PCR (73).

The cell lines were maintained at 37°C in a humidified atmosphere of 5% CO_2_ in culture media with supplements, as detailed in **Supplementary Table 1**. The analyzed cells were treated with drugs (as detailed in **Supplementary Table 2**) the day after cell seeding. For treatments involving nutrient deprivation, the cultivation media was replaced immediately before treatment with DMEM without glucose (11966025, Gibco, Carlsbad, CA, USA) and without additional supplements, i.e., fetal bovine serum and glutamine.

### 4.2. Stable knockdown of DRP1 and AMPKα1

The stable knockdown cells and their respective controls were prepared by lentiviral transduction of constructs encoding target-specific shRNA or scramble shRNA under a constitutive promoter (shDRP1 – sc-43732-V, shAMPKα1 – sc-29673-V, scramble control shCTRL – sc-108080, all from Santa Cruz Biotechnology, Inc., Dallas, TX, USA) according to the manufacturer’s instructions. Subsequently, efficiently transduced cells were selected by two rounds of 5-day treatment with 2.5 µg/ml puromycin (sc-108071A, Santa Cruz Biotechnology). Individual cell clones were isolated by limiting dilution seeding in 96-well plates and exposed to 2.5 µg/ml puromycin every 6th passage.

### 4.3. Cell viability MTT assay

Cells were seeded into 96-well plates at a density of 2000 cells/well. After a 72-hour drug treatment incubation period, an MTT assay was performed. Thiazolyl blue tetrazolium bromide (#M2128, Sigma‒Aldrich, St. Louis, MO, USA) was added to reach a final concentration of 0.455 mg/ml, and the plates were incubated for 3 hours under standard conditions. Subsequently, the medium was aspirated, and the formazan crystals were solubilized with 200 μL of DMSO. The absorbance of each well was measured using a Sunrise Absorbance Reader (Tecan, Männedorf, Switzerland).

To compare drug sensitivity, absolute half-maximal inhibitory concentrations (IC_50_) were determined for individual cell lines and cell clones. They were derived from nonlinear regression of MTT assay datasets, which consisted of absorbance values of individual tested concentrations normalized to those of untreated control cells. Nonlinear regression with variable slope was performed using GraphPad Prism 8.0.2 software (GraphPad Software, San Diego, CA, US). Absolute IC_50_ values were calculated from the nonlinear regression parameters using the following formula: relative IC_50_*(((50-top)/(bottom-50))^(−1/hill slope)).

### 4.4. Cell growth analysis

Cells were seeded in at least 3 technical replicates into 96-well plates at a density of 1500 cells/well. The day after seeding, cell confluency was determined using an IncuCyte® SX1 live-cell imaging system (Sartorius, Göttingen, Germany) every 4 hours over a 72-hour period to monitor cell growth continuity. The cell growth rate was calculated as the ratio of cell confluence in each individual well at the end and beginning of the 72-hour analysis.

### 4.5. Western blotting

Analyzed cells were harvested in RIPA lysis buffer (2 mM EDTA, 1% IGEPAL® CA-630, 0.1% SDS, 8.7 mg/ml sodium chloride, 5 mg/ml sodium deoxycholate, 50 mM Tris-HCl) supplemented with cOmplete™ Mini Protease Inhibitor Cocktail (#11836170001, Roche, Basel, Switzerland) and PhosSTOP (#4906837001, Roche) to prepare whole-cell extracts. A total of 20 µg of protein was denatured using Laemmli sample buffer and heat, loaded into 10% polyacrylamide gels, electrophoretically resolved, and transferred onto PVDF membranes (#1620177, Bio-Rad Laboratories, Hercules, CA, USA). The membranes were then blocked with a solution of 5% nonfat dry milk or bovine serum albumin (#A7906, Sigma‒Aldrich) in Tris-buffered saline supplemented with 0.05% Tween-20 (#93773, Sigma‒Aldrich) for at least 1 hour, followed by overnight incubation at 4°C with primary antibodies on a rocking platform. Next, the membranes were incubated for at least 1 hour at room temperature with secondary HRP-linked antibodies. Details of the antibodies, including dilutions, respective blocking agents and sample heat denaturation temperatures, are provided in **Supplementary Table 3**. Chemiluminescence was detected after a 5-minute incubation with ECL™ Prime Western Blotting Detection Reagent (#RPN2236, Cytiva, Marlborough, MA, USA) using the Azure C600 imaging system (Azure Biosystems, Dublin, CA, USA).

Densitometric image analysis was performed using the gel analysis tool in ImageJ (Fiji) software (NIH, Bethesda, MD, USA), version 2.1.0/1.53c. The detected signal of the protein of interest was normalized to that of the loading control (GAPDH, α-tubulin, or β-actin) provided on the same gel. Original uncropped blots of all replicates are included in the supplementary Original Data file.

### 4.6. Immunostaining

Cells were seeded on glass coverslips. After 2 days of cultivation, the cells were rinsed with PBS, fixed with 3% paraformaldehyde (#158127, Sigma‒Aldrich) for 20 minutes, and permeabilized with 0.2% Triton X-100 (#04807423, MP Biomedicals, Irvine, CA, USA) for 1 minute. The fixed and permeabilized cells were then exposed to a blocking solution (3% bovine serum albumin (#A7906, Sigma‒Aldrich) in PBS) for 20 minutes and incubated with primary and subsequently secondary antibodies diluted in the blocking solution for at least 1 hour at 37°C.

The list of primary and secondary antibodies is provided in **Supplementary Table 3**. Nuclei were stained with 1 μg/ml Hoechst 33342 (#H1399, Invitrogen, Carlsbad, CA, USA). The coverslips were mounted using ProLong™ Diamond Antifade (#P36961, Invitrogen). Z-stacked images were captured using a Leica SP8 confocal microscope (Leica, Wetzlar, Germany) and processed as maximum intensity projections using LAS X software (Leica, 3.4.218368).

### 4.7. Mitochondrial network morphology analysis

The ImageJ (Fiji) plug-in tool Mitochondrial Analyzer (74) was used to analyze the mitochondrial network morphology on a per-field-of-view basis from 2D maximum intensity projections of z-stack confocal images of the TOMM20 fluorescence channel. The preprocessing parameters were set as follows: rolling (microns), 1; radius, 2; max slope, 2; and gamma, 0.8. The applied commands included subtracting the background, sigma filter plus, enhancing local contrast, and adjusting gamma. The threshold method used was the weighted mean method with a block size of 1.75 microns and a c value of 1.25. Postprocessing commands included despeckling and removing outliers with a 2-pixel radius. At least 2000 mitochondria in at least 4 fields of view were analyzed for each biological replicate.

### 4.8. Gene expression transcriptomic analysis

The R2: Genomics Analysis and Visualization Platform (http://r2.amc.nl) was used for single-gene survival analysis. The expression of the mitochondrial fission mediator DRP1-encoding gene *DNM1L1* was utilized to group patients, for whom Kaplan‒Meier survival curves were generated from publicly available osteosarcoma (Kuijjer) (75) and rhabdomyosarcoma (Williamson) (76) transcriptomic datasets. Kaplan–Meier curves were plotted for the high (blue) and low (red) expression groups established by the scan cutoff mode with a minimal group size of 20 patients. Additionally, the p value was calculated from the plotted curves using a two-sided log-rank test.

### 4.9. Statistical analysis

All experiments were performed with at least three independent biological replicates; further details are provided in the figure legends. Line graphs and bar graphs display the mean ± standard deviation (SD). In the bar graphs, individual data points represent independent biological replicates, except for the general comparison of the control and knockdown cell clone groups in Figs. 5a, e; 6; 7; 8a, g; S17c; S18d; S20b, where individual data points refer to the mean value determined for individual cell clones from at least three independent biological replicates. Further exceptions are Figs. 5c, d, f, g; 8c, d; S9; S16, where individual data points represent the IC_50_ determined for each individual cell clone. Statistical analysis of Kaplan‒Meier survival curves was performed using the R2: Genomics Analysis and Visualization Platform (http://r2.amc.nl) with a two-sided log-rank test; otherwise, statistical analysis was conducted using GraphPad Prism 8.0.2. software. An unpaired two-tailed Student’s t-test was applied when comparing two groups; otherwise, one-way ANOVA followed by Tukey’s multiple comparison test was used, assuming a normal data distribution and similar variance between the compared groups. p values < 0.05 were considered to indicate statistical significance; *p<0.05, **p<0.01, ***p<0.001, #p<0.0001; ns refers to a p value ≥0.05.

## DATA AVAILABILITY

All data are included in the article and supporting information. Additional supporting data are available from the corresponding author upon reasonable request.

## ACKNOWLEDGMENTS

This work was supported by the Czech Science Foundation (No. GJ20-00987Y). JS and KB acknowledge the project National Institute for Cancer Research (Programme EXCELES, ID Project No. LX22NPO5102)— Funded by the European Union—Next Generation EU. The authors thank to Johana Marešová and Dagmar Štodtová for their skilled technical assistance.

## AUTHOR CONTRIBUTIONS

**Karolina Borankova:** Conceptualization, Writing - Original Draft, Methodology, Investigation, Supervision, Formal analysis, Data Curation, Visualization, Writing - Review & Editing. **Matyas Solny:** Investigation, Formal analysis. **Maria Krchniakova:** Investigation, Formal analysis, Writing - Review & Editing. **Jan Skoda:** Conceptualization, Writing - Original Draft, Methodology, Resources, Supervision, Visualization, Project administration, Funding acquisition, Writing - Review & Editing.

In addition to the CRediT authorship contributions listed above, the detailed contributions are as follows:

KB and JS conceptualized and drafted the manuscript; KB, MK, and JS designed the experiments; KB conducted key experiments in sarcoma cells; MS performed immunoblotting analysis of RD cells treated with doxorubicin or cisplatin and tested the sensitivity of rhabdomyosarcoma cells to bafilomycin/rapamycin alone or in combination with chemotherapy; MK performed immunoblotting analysis of osteosarcoma cells treated with cisplatin; KB performed the statistical analyses; KB, MS, MK and JS analyzed the data; and KB, MK and JS wrote and edited the manuscript. All the authors have read and approved the final manuscript.

## SUPPLEMENTARY MATERIAL

### SUPPLEMENTARY FIGURES

**Fig. S1:**
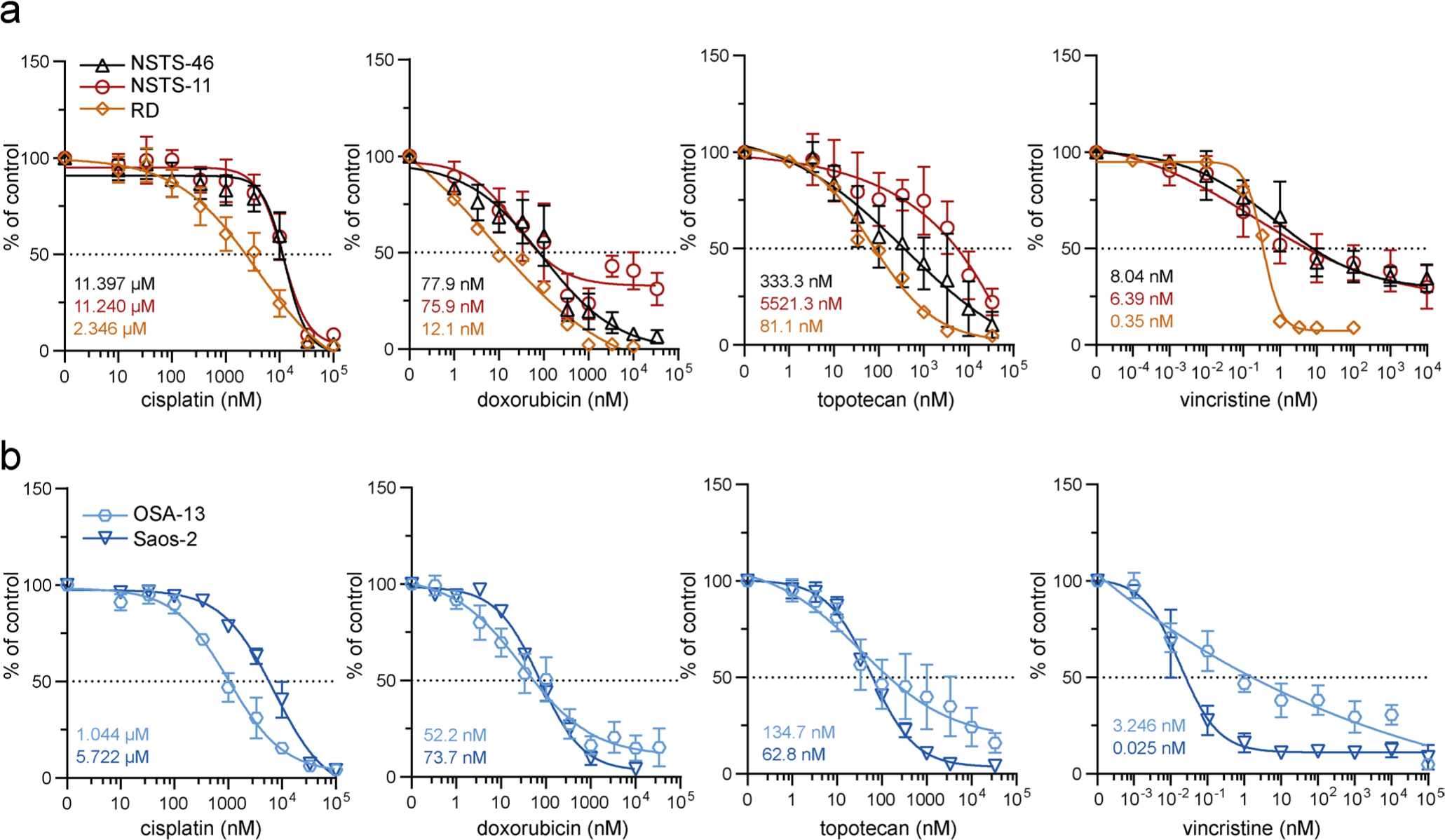
Chemotherapy dose‒response curves showing the IC50 values in sarcoma cells. **(a, b)** MTT viability assay analysis of rhabdomyosarcoma (a) and osteosarcoma (b) cells exposed to the indicated concentrations of chemotherapy drugs for 72 hours was used to establish the IC50 values. The data points represent the mean ± SD, biological n≥3, technical n=3.

**Fig. S2:**
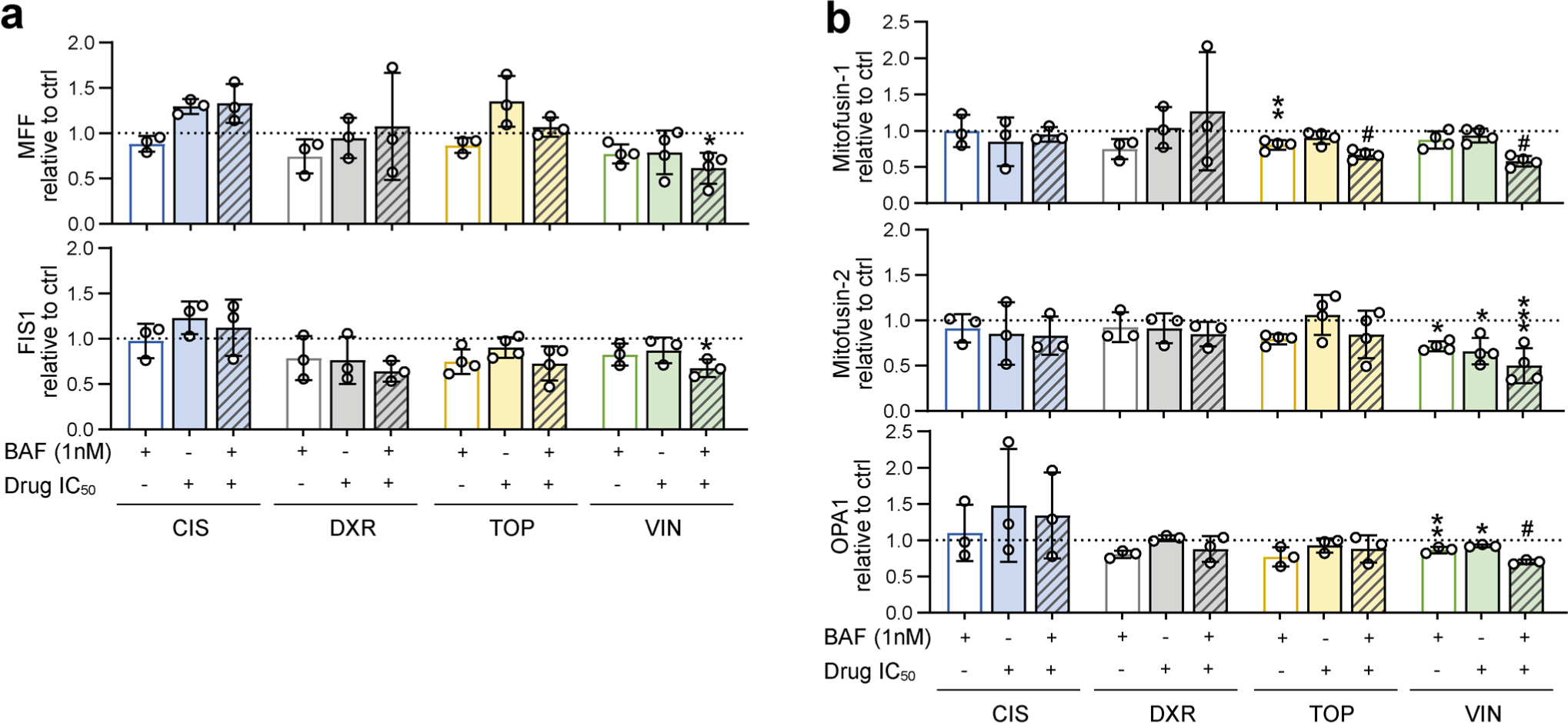
Extended densitometric analysis of mitochondrial fission and fusion-related proteins detected in rhabdomyosarcoma RD cells exposed to bafilomycin A1 and/or chemotherapy drugs. **(a, b)** Densitometric analysis of (a) mitochondrial fission and (b) fusion-related proteins in rhabdomyosarcoma RD cells exposed to 1 nM bafilomycin A1 (BAF), chemotherapy drugs at the IC50 dose, or combinations of BAF and chemotherapy drugs for 72 hours. Normalized protein levels are plotted relative to the untreated controls as the mean ± SD. BAF – bafilomycin A1, CIS – cisplatin, DXR – doxorubicin, TOP – topotecan, VIN – vincristine. Statistical significance was determined by one-way ANOVA followed by Tukey’s multiple comparisons test, *p<0.05, **p<0.01, ***p<0.001, #p<0.0001.

**Fig. S3:**
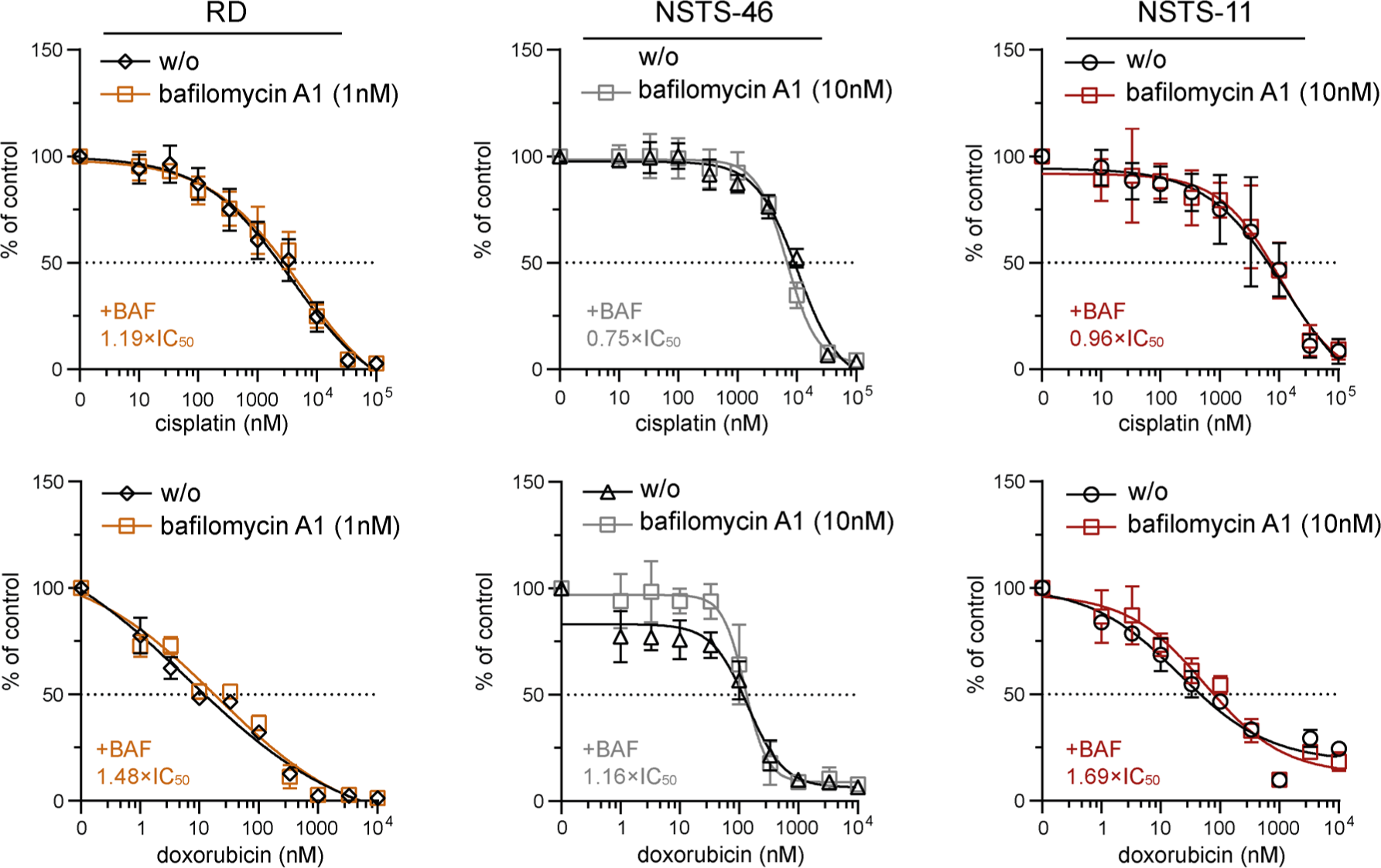
Effect of the autophagy inhibitor bafilomycin A1 on sensitivity to cisplatin and doxorubicin in both bafilomycin A1-sensitive and bafilomycin A1-resistant rhabdomyosarcoma cells. MTT viability assay analysis of bafilomycin A1-sensitive (RD) and bafilomycin A1-resistant (NSTS-46, NSTS-11) rhabdomyosarcoma cells exposed to the indicated concentrations of cisplatin and doxorubicin alone or in the presence of a sublethal bafilomycin dose (RD – 1 nM; NSTS-46, NSTS-11 – 10 nM) for 72 hours revealed that bafilomycin A1 did not significantly increase sensitivity to cisplatin or doxorubicin. The bafilomycin A1-induced IC50 fold change is indicated as the ratio of the IC50 determined for the combination of bafilomycin A1 and chemotherapy drugs to the IC50 determined for chemotherapy drugs alone. The data points represent the mean ± SD, biological n≥3, technical n=3.

**Fig. S4:**
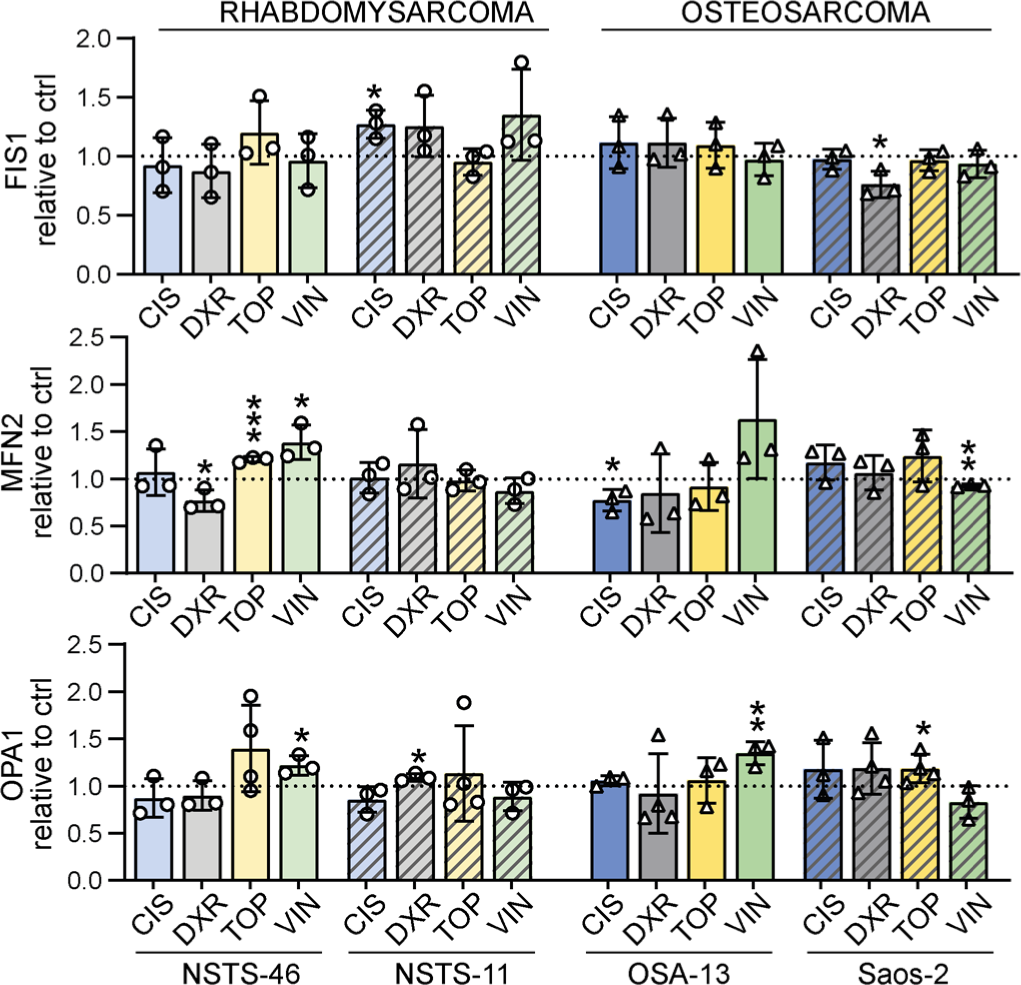
Extended densitometric analysis of mitochondrial fission and fusion-related proteins detected in sarcoma cells exposed to chemotherapy. Densitometric analysis of Western blotting detection of mitochondrial fission and fusion-related proteins in both rhabdomyosarcoma (NSTS-46 and NSTS-11) and osteosarcoma (OSA-13 and Saos-2) cells exposed to chemotherapy drug IC50 for 72 hours. Normalized protein levels are plotted relative to the untreated controls as the mean ± SD. CIS – cisplatin, DXR – doxorubicin, TOP – topotecan, VIN – vincristine. Statistical significance was determined by unpaired two-tailed Student’s t-test, *p<0.05, **p<0.01, ***p<0.001.

**Fig. S5:**
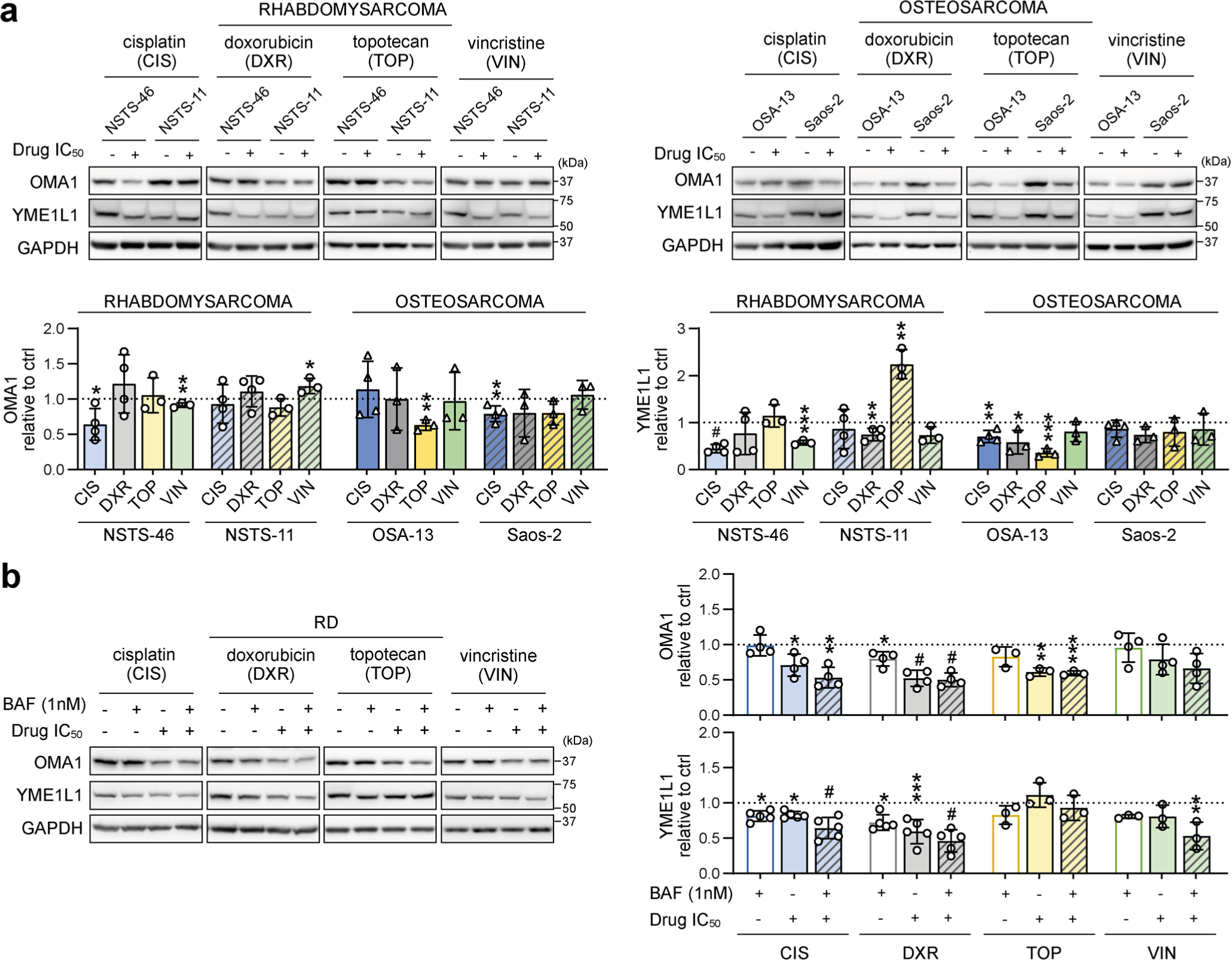
Chemotherapy exposure predominantly downregulated mitochondrial stress sensors in sarcoma cells. **(a)** Western blotting and densitometric analysis showed that the mitochondrial stress sensors OMA1 and YME1L1 were predominantly downregulated in rhabdomyosarcoma and osteosarcoma cells exposed to IC50 chemotherapy for 72 hours. Normalized protein levels are plotted relative to the untreated controls as the mean ± SD. **(b)** Western blotting and densitometric analysis of mitochondrial stress sensors in RD rhabdomyosarcoma cells after 72 hours of exposure to 1 nM bafilomycin A1 (BAF), chemotherapy drugs at the IC50 dose, or combinations of BAF and chemotherapy drugs revealed that OMA1 and YME1L1 are downregulated by chemotherapy both in the absence and presence of bafilomycin A1-mediated autophagy inhibition. Normalized protein levels are plotted relative to the untreated controls as the mean ± SD. BAF – bafilomycin A1, CIS – cisplatin, DXR – doxorubicin, TOP – topotecan, VIN – vincristine. Statistical significance was determined by unpaired two-tailed Student’s t-test (a) and by one-way ANOVA followed by Tukey’s multiple comparisons test (b), *p<0.05, **p<0.01, ***p<0.001, #p<0.0001.

**Fig. S6:**
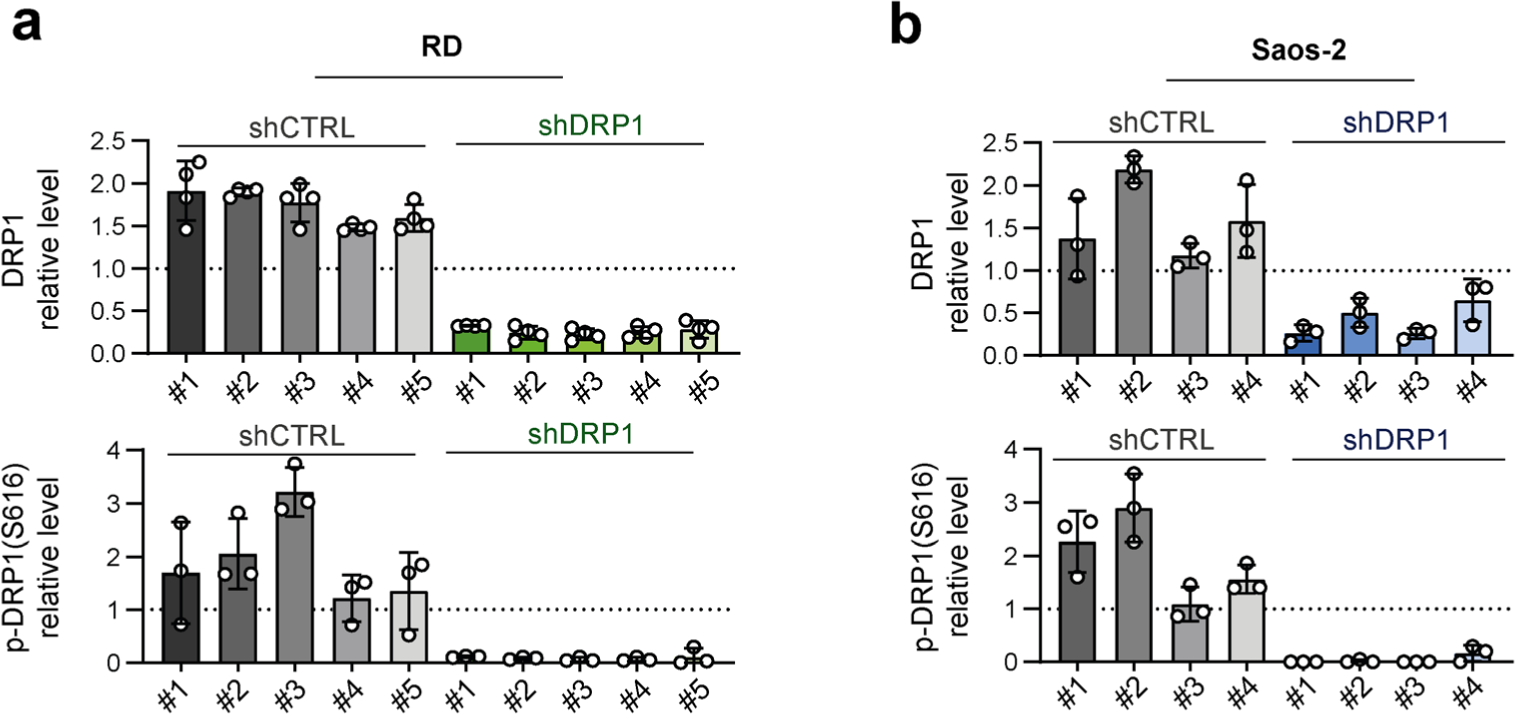
Individual cell clone analysis confirmed the efficiency of stable DRP1 downregulation in sarcoma cells. **(a, b)** Densitometric analysis of Western blotting of DRP1 in individual rhabdomyosarcoma RD (a) and osteosarcoma Saos-2 (b) control (shCTRL) and DRP1-downregulated (shDRP1) cell clones showed that significant DRP1 downregulation resulted in a marked decrease in its activating form, p-DRP1(S616). Normalized protein levels detected in individual clones are plotted relative to the average level as the mean ± SD.

**Fig. S7:**
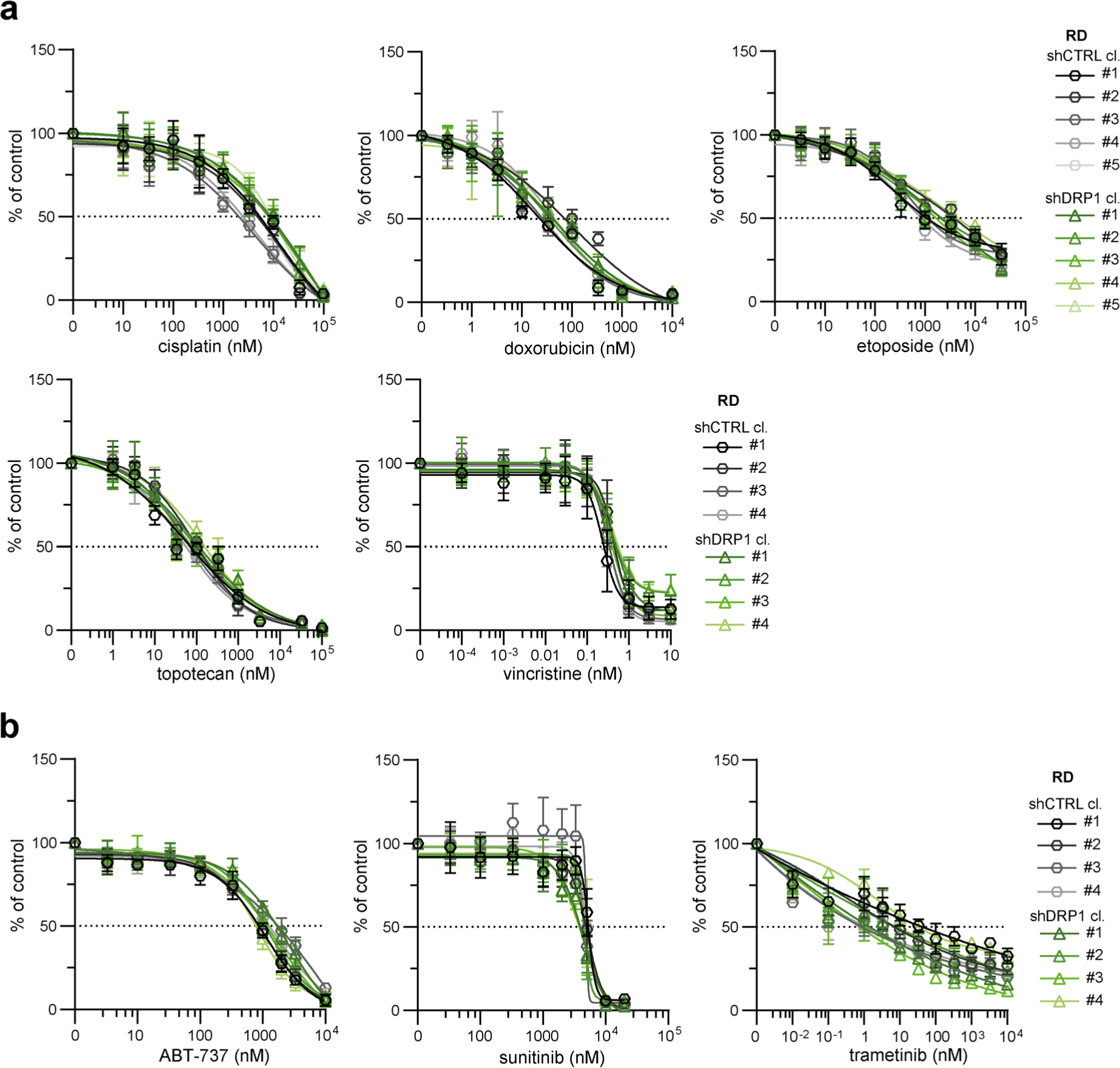
Stable DRP1 downregulation did not mitigate drug resistance in RD rhabdomyosarcoma cells. **(a, b)** MTT viability assay analysis of rhabdomyosarcoma RD control (shCTRL) and DRP1-downregulated (shDRP1) cell clones exposed to standard chemotherapy drugs (a) or targeted inhibitors (b) at the indicated concentrations for 72 hours showed that DRP1 knockdown did not attenuate drug resistance. The data points represent the mean ± SD, biological n≥3, technical n=3.

**Fig. S8:**
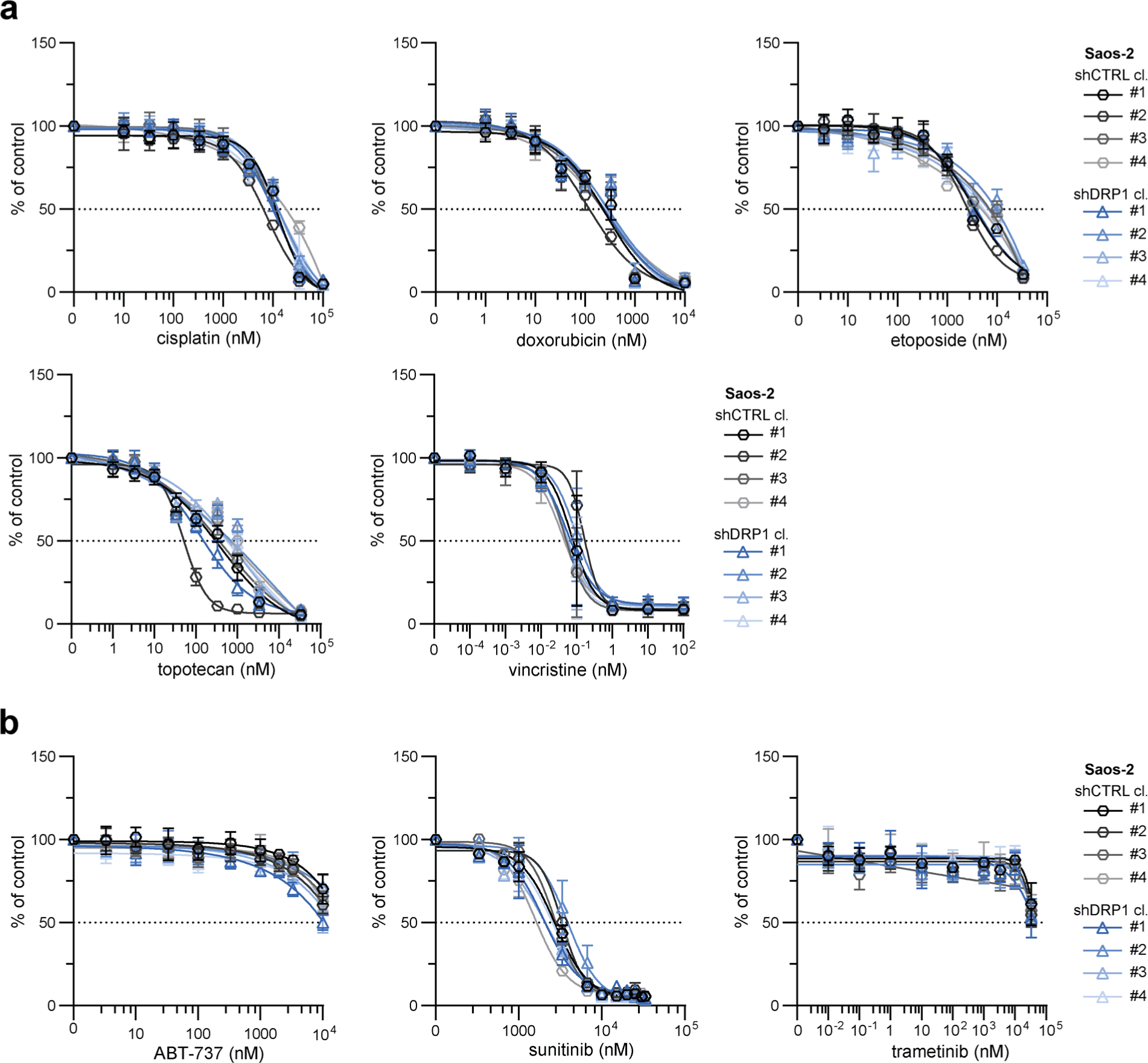
Stable DRP1 downregulation did not mitigate drug resistance in Saos-2 osteosarcoma cells. **(a, b)** MTT viability assay analysis of Saos-2 control (shCTRL) and DRP1-downregulated (shDRP1) osteosarcoma cell clones exposed to standard chemotherapy drugs (a) or targeted inhibitors (b) at the indicated concentrations for 72 hours showed that DRP1 knockdown did not attenuate drug resistance. The data points represent the mean ± SD, biological n≥3, technical n=3.

**Fig. S9:**
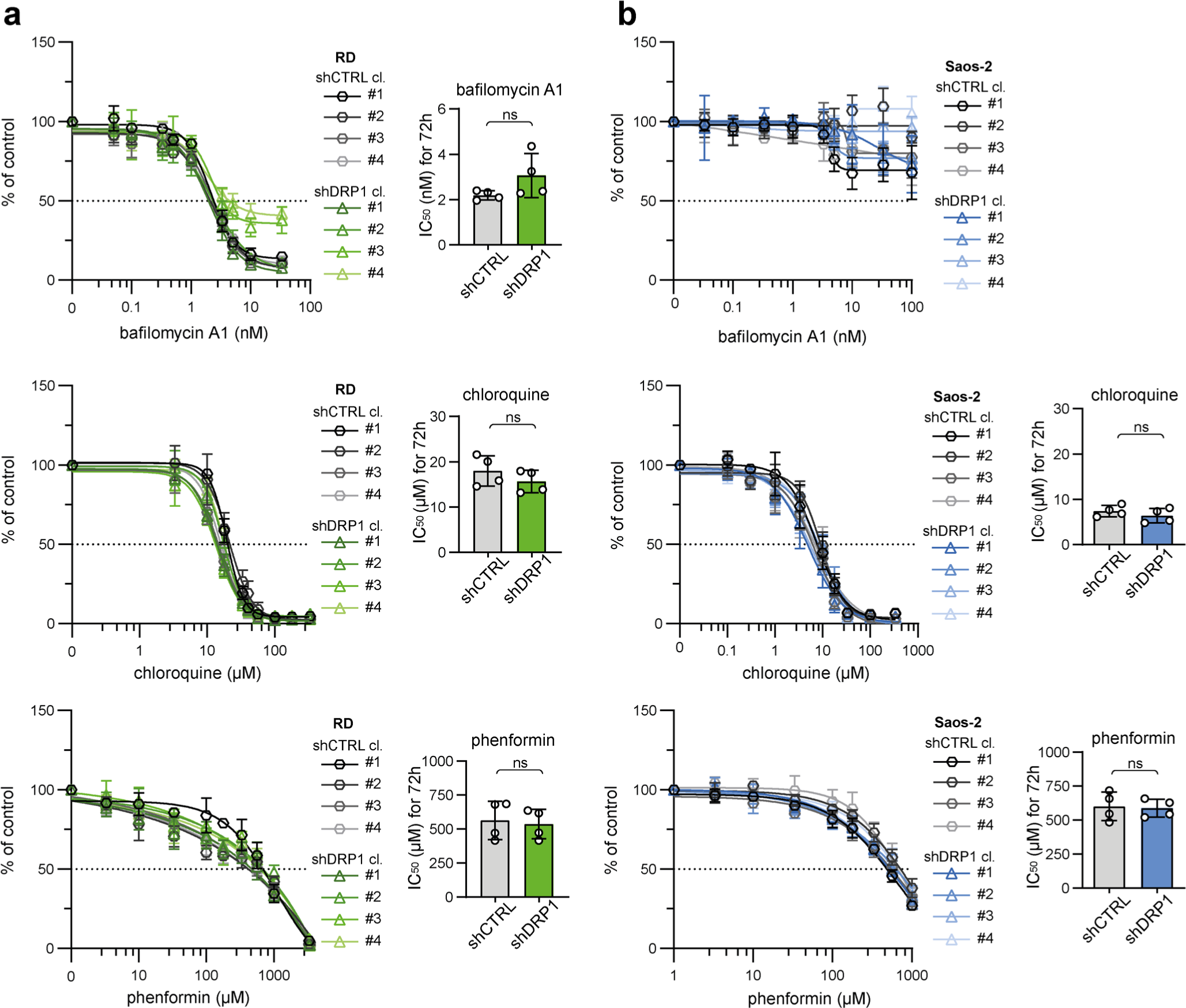
Stable DRP1 downregulation did not affect sarcoma cell sensitivity to autophagy or respiratory complex I inhibition. **(a, b)** MTT viability assay analysis of rhabdomyosarcoma RD (a) and osteosarcoma Saos-2 (b) control (shCTRL) and DRP1-downregulated (shDRP1) cell clones exposed to autophagy inhibitors (bafilomycin A1 and chloroquine) and a respiratory complex I inhibitor (phenformin) at the indicated concentrations for 72 hours showed that sensitivity to these inhibitors was not affected by DRP1 downregulation. The data points represent the mean ± SD, biological n≥3, technical n=3. The impact of DRP1 downregulation on the sensitivity to these specific inhibitors was assessed by comparison of the IC50 values. The data are plotted as the mean ± SD of the IC50 values determined for individual shCTRL and shDRP1 cell clones. Statistical significance was determined by unpaired two-tailed Student’s t-test, ns – not significant.

**Fig. S10:**
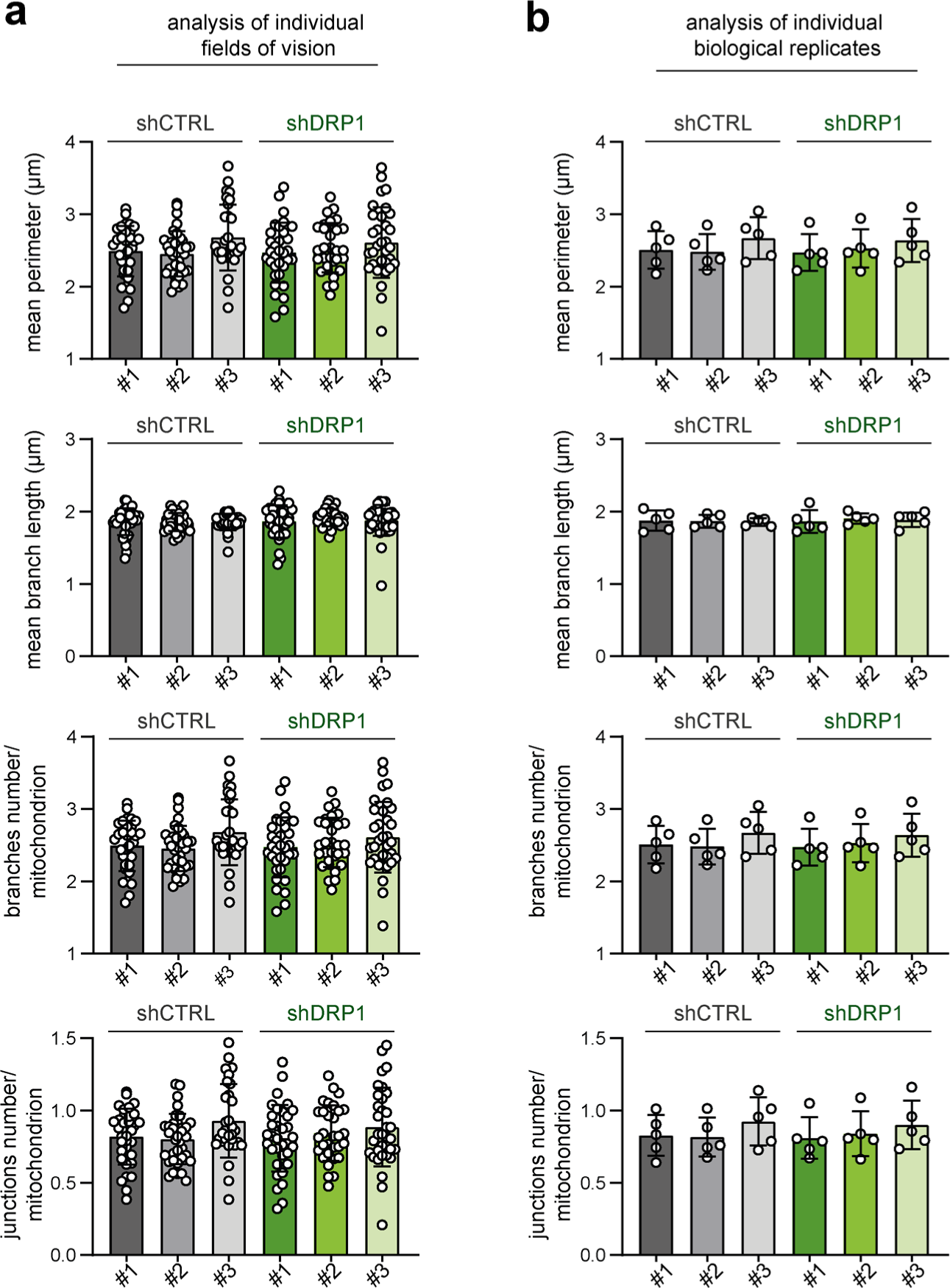
Stable DRP1 downregulation did not affect the mitochondrial network morphology in RD rhabdomyosarcoma cells. **(a,b)** Image analysis using the ImageJ plug-in tool Mitochondria Analyzer showed that DRP1 downregulation did not affect parameters describing the mitochondrial network morphology in rhabdomyosarcoma RD cells. Analysis of both individual fields of view (a) and biological replicates (b) revealed that the mean mitochondrial perimeter, mean mitochondrial branch length, number of mitochondrial branches per mitochondrion, and number of mitochondrial junctions per mitochondrion were not different between individual DRP1-downregulated (shDRP1) clones and their control counterparts (shCTRL).

**Fig. S11:**
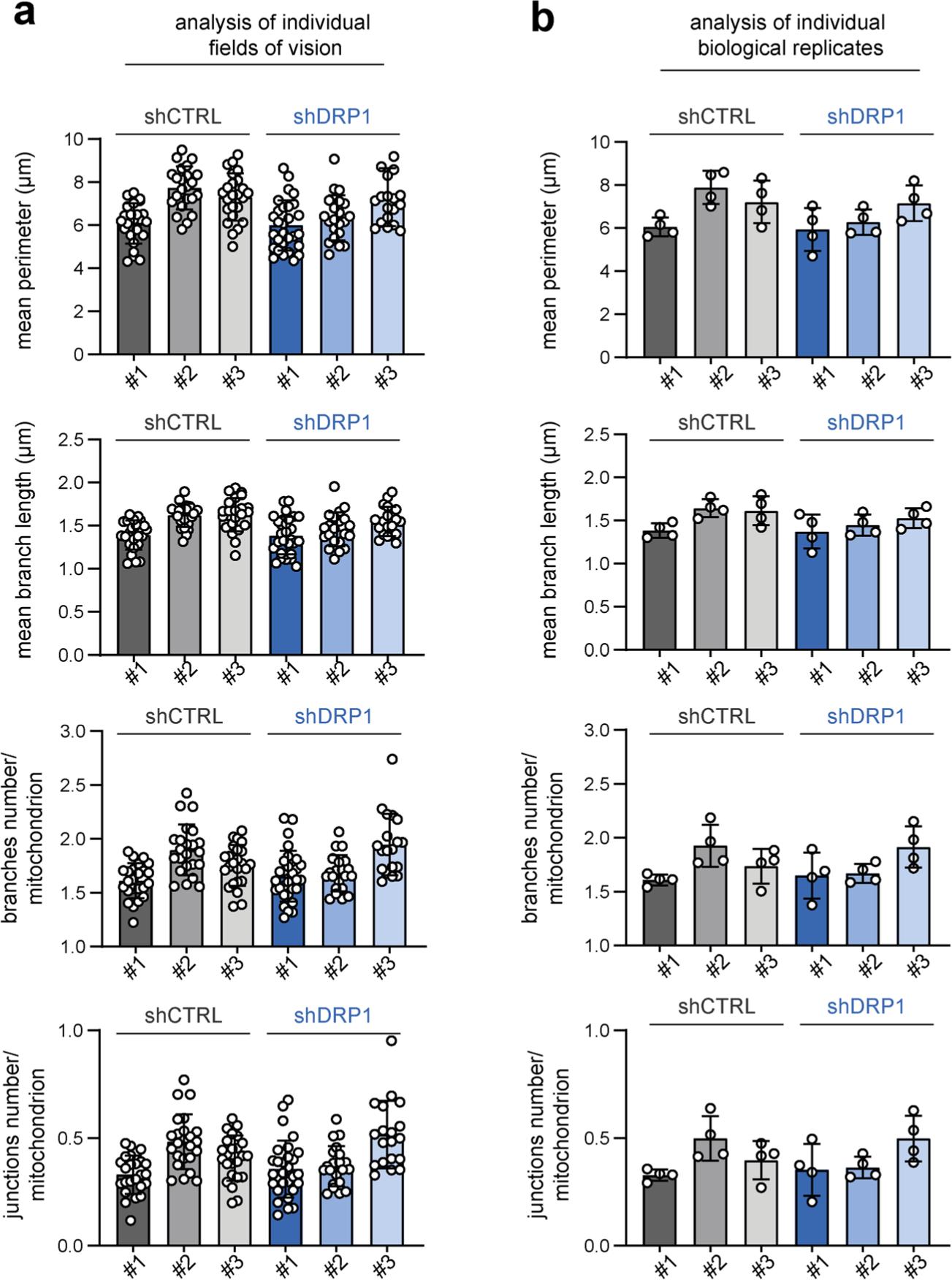
Stable DRP1 downregulation did not affect the mitochondrial network morphology in Saos-2 osteosarcoma cells. **(a, b)** Image analysis using the ImageJ plug-in tool Mitochondria Analyzer showed that DRP1 downregulation did not affect parameters describing the mitochondrial network morphology in Saos-2 osteosarcoma cells. Analysis of both individual fields of view (a) and biological replicates (b) revealed that the mean mitochondrial perimeter, mean mitochondrial branch length, number of mitochondrial branches per mitochondrion, and number of mitochondrial junctions per mitochondrion were not different between individual DRP1-downregulated (shDRP1) clones and their control counterparts (shCTRL).

**Fig. S12:**
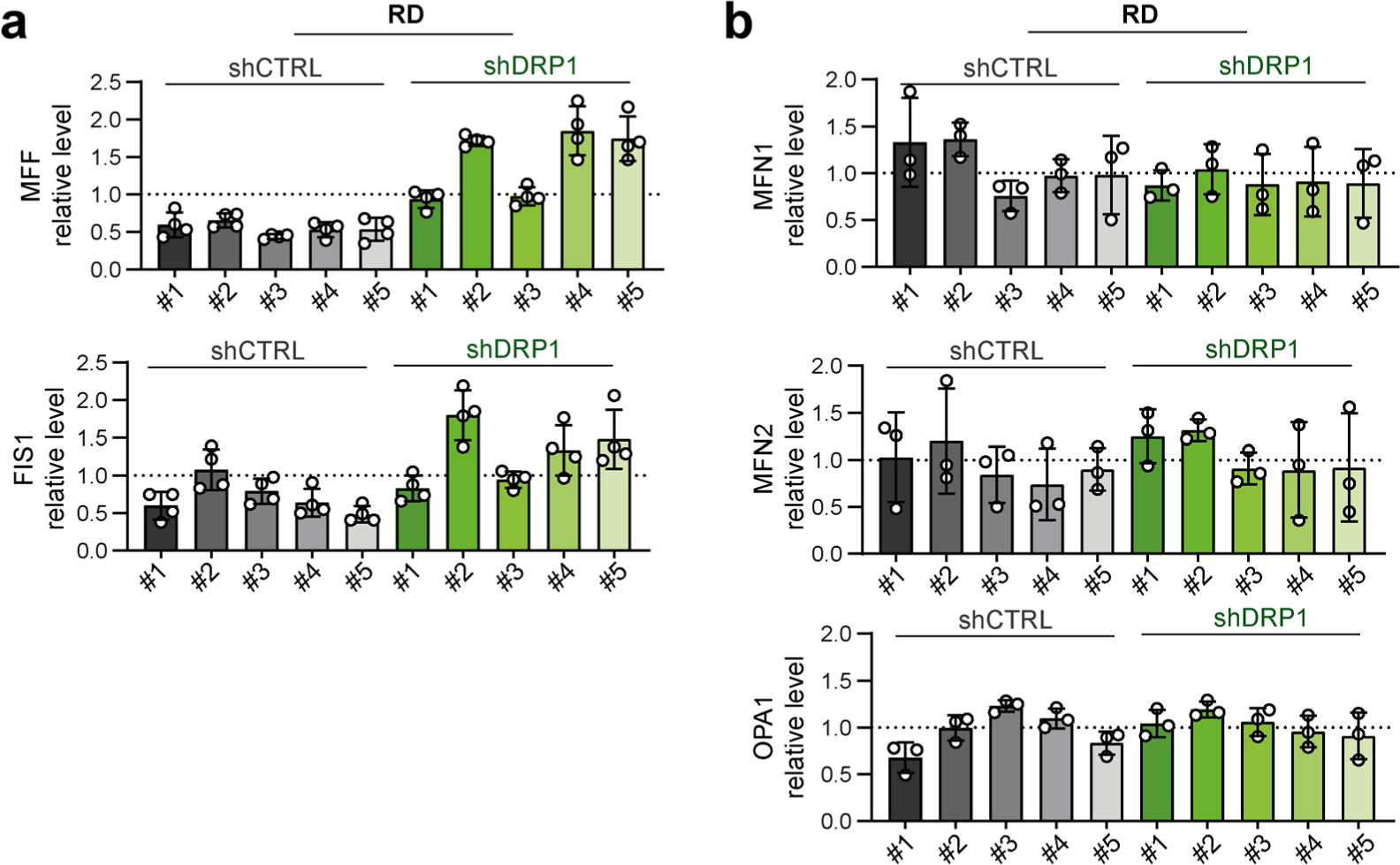
Stable knockdown of the mitochondrial fission mediator DRP1 upregulated the expression of the adaptor proteins MFF and FIS1 in RD rhabdomyosarcoma cells. **(a, b)** Densitometric analysis of Western blotting (Fig. 6b) of mitochondrial fission-(a) and fusion-related (b) proteins in individual rhabdomyosarcoma RD control (shCTRL) and DRP1-downregulated (shDRP1) cell clones showed that DRP1 knockdown resulted in an increase in the levels of DRP1 adaptors MFF and FIS1. Normalized protein levels detected in individual clones are plotted relative to the average level as the mean ± SD.

**Fig. S13:**
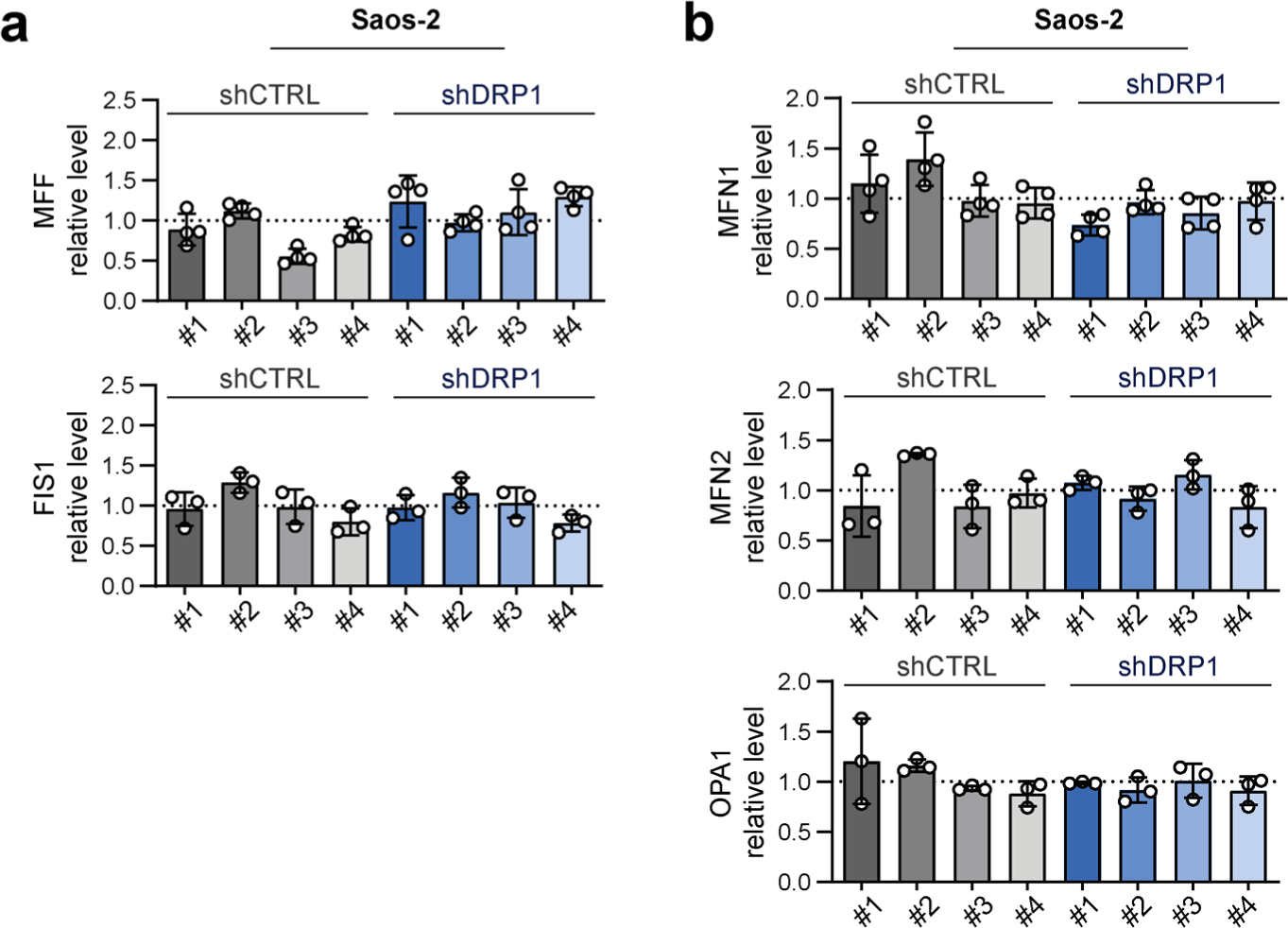
Stable knockdown of the mitochondrial fission mediator DRP1 did not impact the levels of other mitochondrial fission- or fusion-related proteins in Saos-2 osteosarcoma cells. **(a, b)** Densitometric analysis of Western blotting (Fig. 7b) of mitochondrial fission-(a) and fusion-related (b) proteins revealed no significant differences between individual Saos-2 control (shCTRL) and DRP1-downregulated (shDRP1) osteosarcoma cell clones. Normalized protein levels detected in individual clones are plotted relative to the average level as the mean ± SD.

**Fig. S14:**
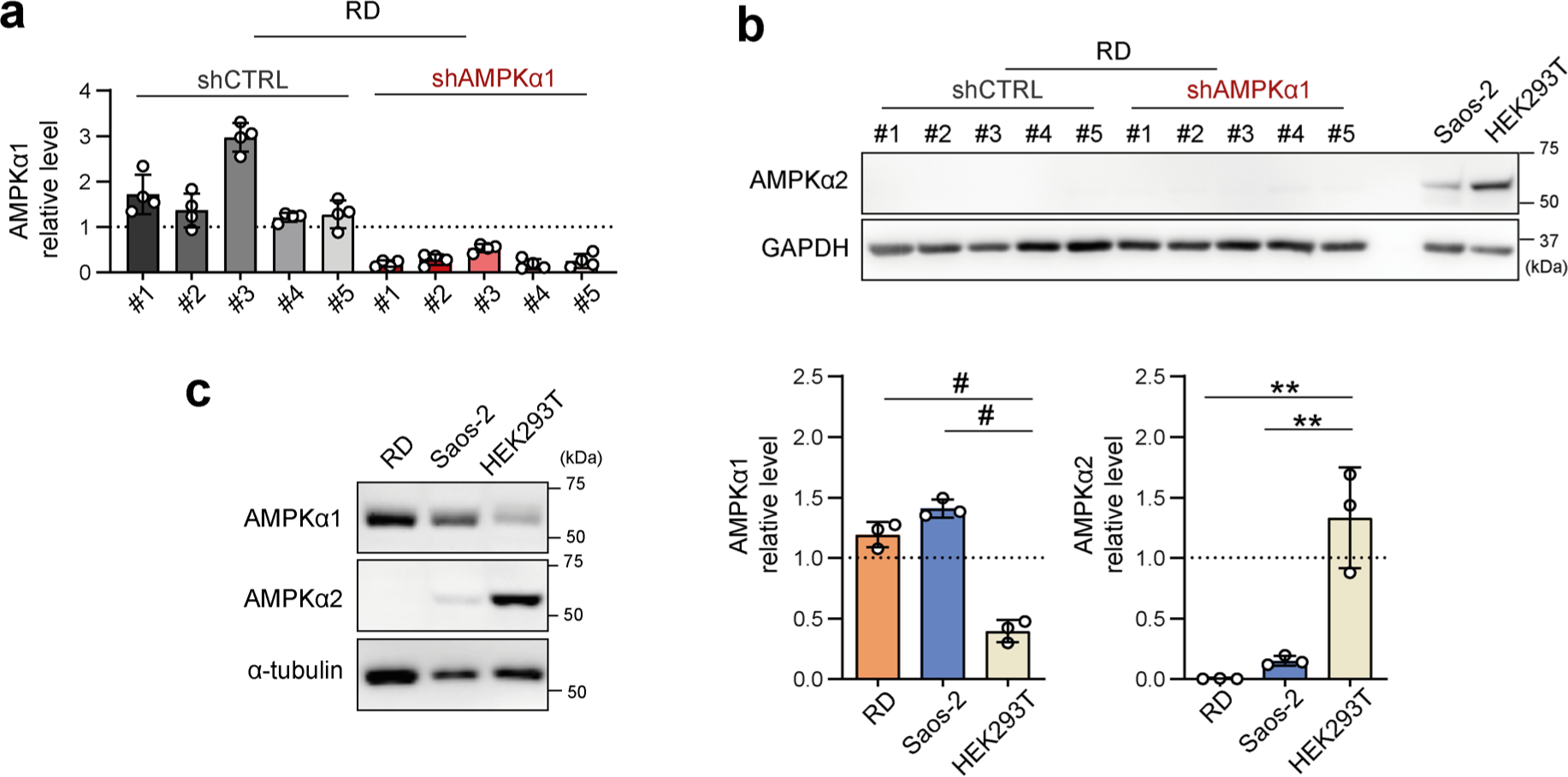
The efficiency of stable AMPKα1 downregulation was confirmed in individual rhabdomyosarcoma RD cell clones, while AMPKα2 expression remained undetectable in all clones. **(a)** Densitometric analysis of Western blotting of AMPKα1 in rhabdomyosarcoma control (shCTRL) and AMPKα1-downregulated (shAMPKα1) cell clones revealed significant downregulation of AMPKα1 in all individual shAMPKα1 clones compared to their control counterparts. Normalized protein levels detected in individual clones are plotted relative to the average level as the mean ± SD. **(b)** Western blot analysis showed that AMPKα2 was undetectable in both the RD shCTRL and shAMPKα1 clones. Saos-2 and HEK293T cells served as positive controls for the detection. **(c)** Western blotting and densitometric analysis of AMPKα1 and AMPKα2 in RD, Saos-2 and HEK293T cells revealed that AMPKα2 expression was undetectable in RD cells, while AMPKα1 expression in RD cells was increased compared with that in HEK293T cells, which show enhanced expression of AMPKα2. Overall, low AMPKα2 expression in Saos-2 cells and undetectable AMPKα2 expression in RD tissues suggest that the AMPKα1 isoform is the key isoform of AMPKα in sarcoma cells. Statistical significance was determined by one-way ANOVA followed by Tukey’s multiple comparisons test (c), **p<0.01, #p<0.0001.

**Fig. S15:**
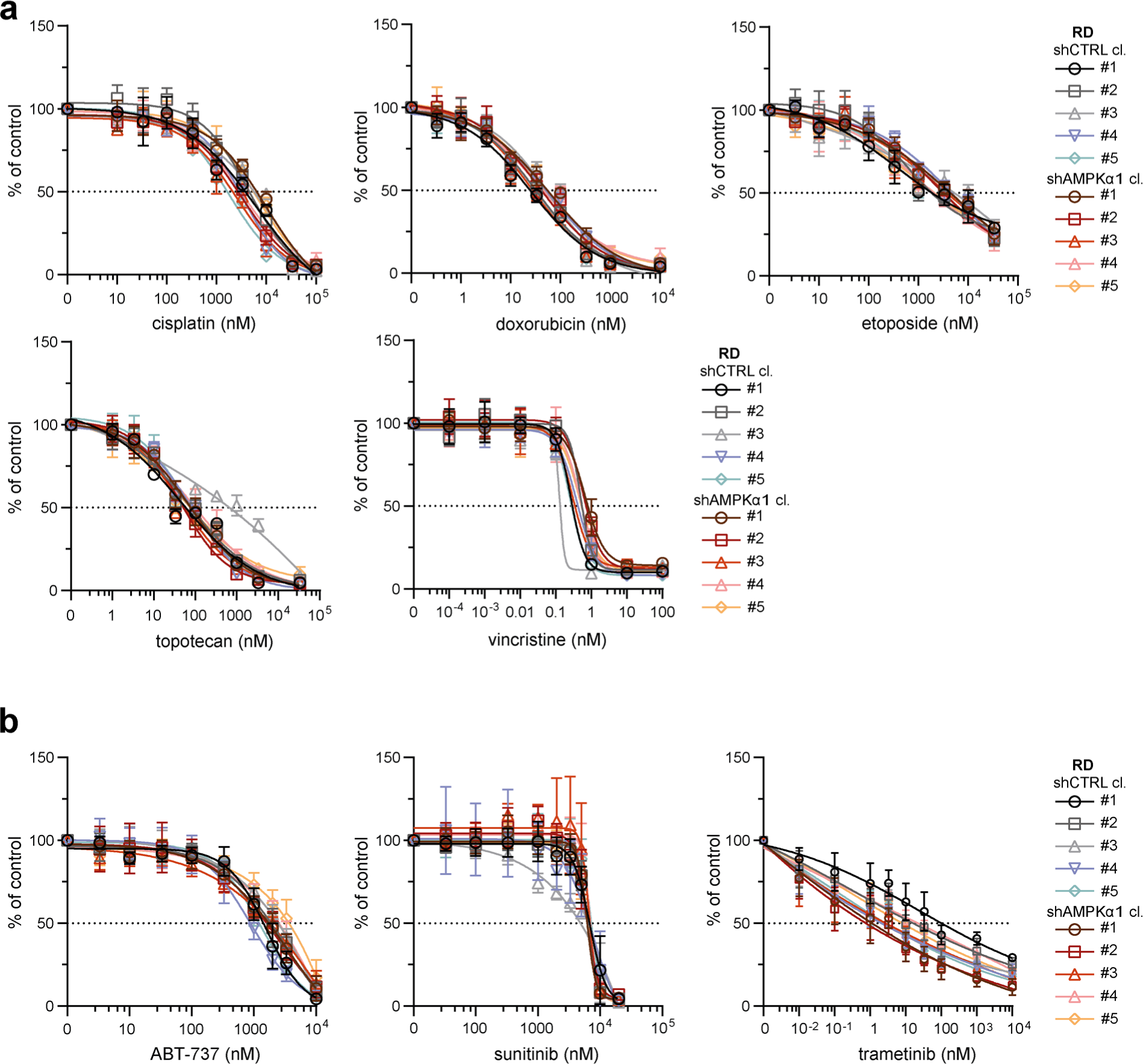
Stable AMPKα1 downregulation did not mitigate drug resistance in RD rhabdomyosarcoma cells. **(a, b)** MTT viability assay analysis of rhabdomyosarcoma RD control (shCTRL) and AMPKα1-downregulated (shAMPKα1) cell clones exposed to standard chemotherapy drugs (a) or targeted inhibitors (b) at the indicated concentrations for 72 hours showed that AMPKα1 knockdown did not attenuate drug resistance. The data points represent the mean ± SD, biological n≥3, technical n=3.

**Fig. S16:**
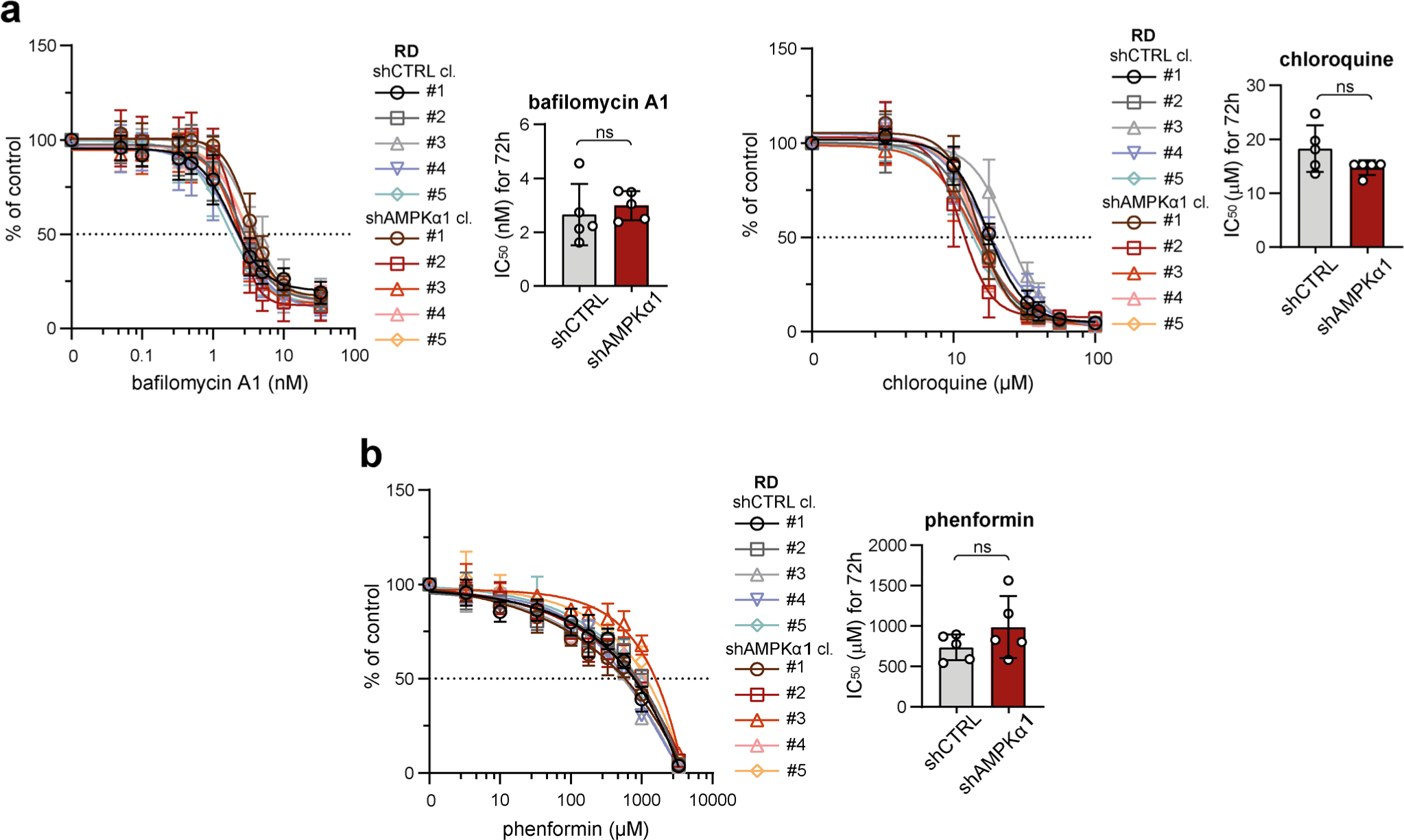
Stable AMPKα1 downregulation did not affect rhabdomyosarcoma cell sensitivity to autophagy or respiratory complex I inhibition. **(a, b)** MTT viability assay analysis of rhabdomyosarcoma RD control (shCTRL) and AMPKα1-downregulated (shAMPKα1) cell clones exposed to autophagy inhibitors (bafilomycin A1, chloroquine) and a respiratory complex I inhibitor (phenformin) at the indicated concentrations for 72 hours showed that sensitivity to these inhibitors was not affected by AMPKα1 downregulation. The data points represent the mean ± SDs, biological n≥3, technical n=3. The resistance of control and AMPKα1-downregulated RD cells to these specific inhibitors was compared using drug IC50 values. The data are plotted as the mean ± SDs of the IC50 values determined for individual shCTRL and shAMPKα1 cell clones. Statistical significance was determined by unpaired two-tailed Student’s t-test, ns – not significant.

**Fig. S17:**
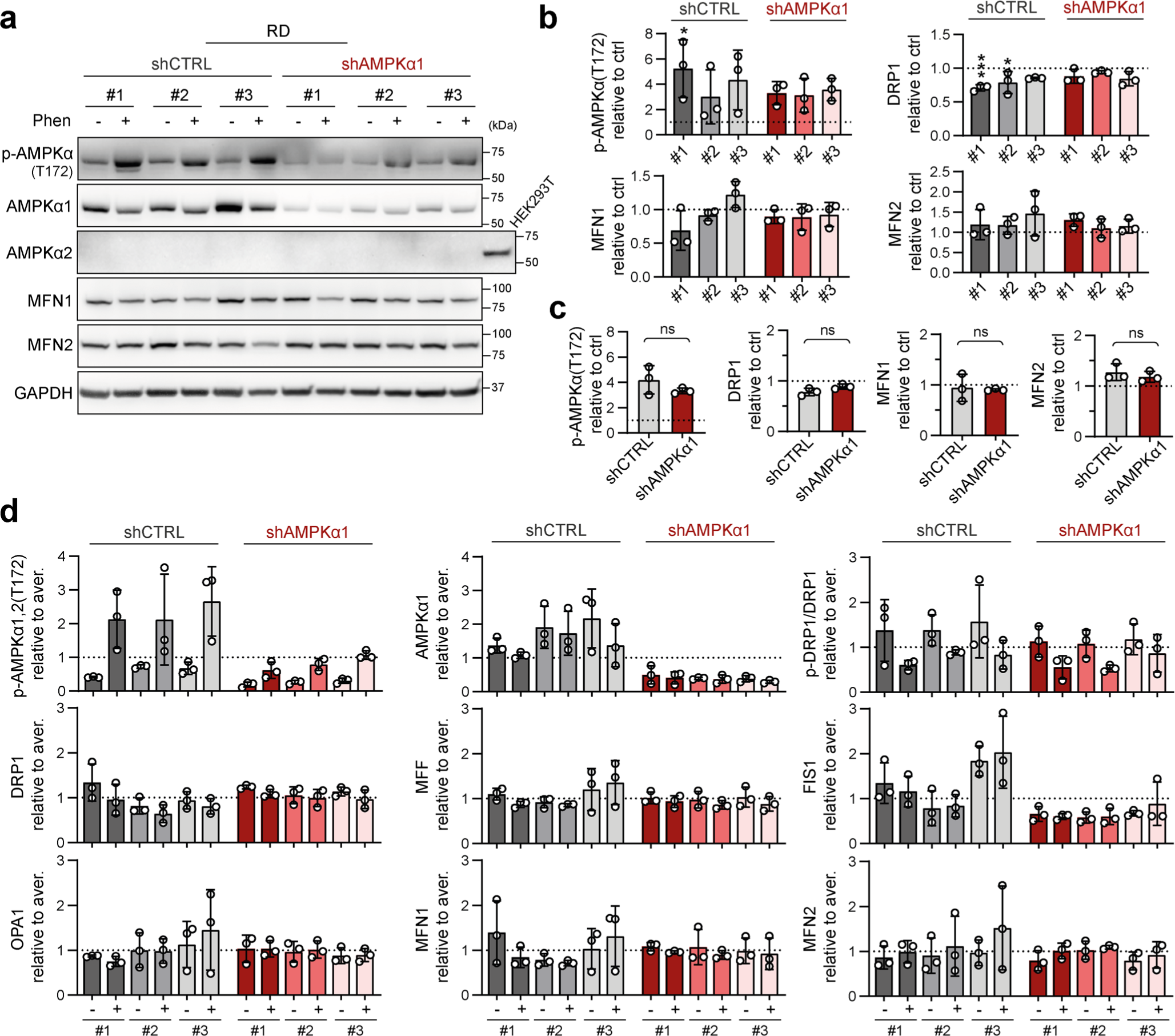
Stable AMPKα1 downregulation did not affect stress-induced modulation of the expression of mitochondrial fission or fusion-related proteins in rhabdomyosarcoma RD cells. **(a-d)** Extension of **Fig. 8e-g**: Western blotting (a) and densitometric analysis (b-d) of rhabdomyosarcoma RD shCTRL and shAMPKα1 cell clones exposed to 1 mM phenformin for 72 hours did not reveal any significant impact of AMPKα1 knockdown on stress-induced modulation of mitochondrial fission and fusion-related proteins. **(b)** Normalized protein levels detected in individual cell clones are plotted relative to the untreated controls as the mean ± SD. **(c)** Comparison of the phenformin-induced modulation of mitochondrial fission and fusion-related proteins detected in the RD shCTRL and shAMPKα1 clone groups. The mean normalized protein levels relative to the untreated controls are plotted as the mean ± SD. **(d)** To display the complete variability of shCTRL and shAMPKα1 cell clones in response to phenformin treatment, normalized protein levels detected in both untreated and phenformin-treated individual cell clones are plotted relative to the average level as the mean ± SD. Statistical significance was determined by one-way ANOVA followed by Tukey’s multiple comparisons test (b) and by unpaired two-tailed Student’s t-test (c), *p<0.05, **p<0.01, ns – not significant.

**Fig. S18:**
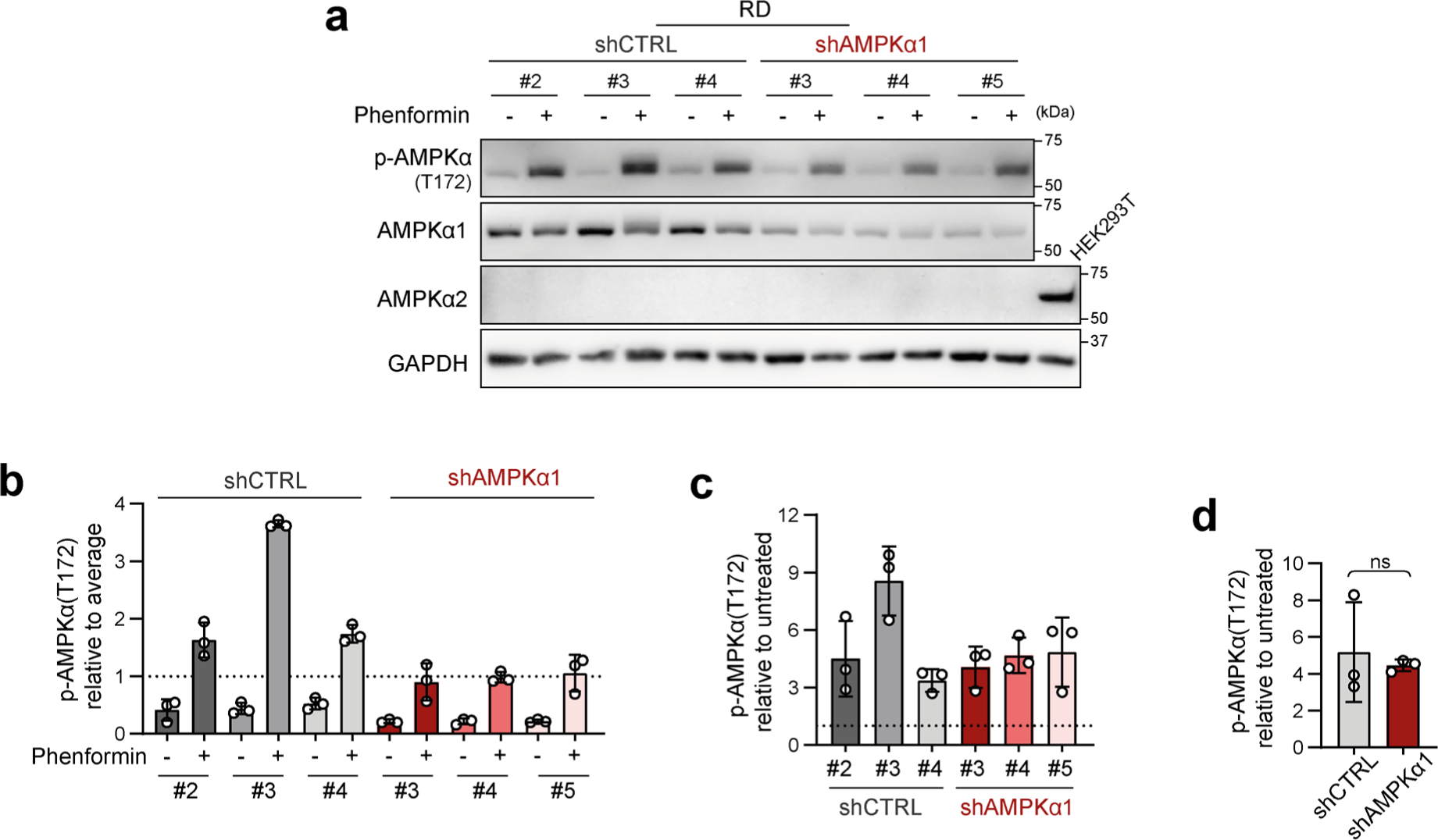
Rhabdomyosarcoma AMPKα1-knockdown cell clones preserve their AMPK signaling capacity independent of AMPKα2. **(a-d)** Western blotting (a) and densitometric analysis (b-d) of AMPKα1 and its activated form p-AMPKα1,2(T172) in rhabdomyosarcoma RD control (shCTRL) and AMPKα1-downregulated (shAMPKα1) cell clones exposed to 5 mM phenformin for 90 minutes upon nutrient deprivation due to glucose, glutamine and serum depletion. **(b)** Normalized p-AMPKα1,2(T172) levels detected in individual cell clones, both untreated and phenformin treated, are plotted relative to the average level as the mean ± SD. Phenformin-treated shAMPKα1 cell clones showed slightly limited upregulation of p-AMPKα1,2(T172), but these levels remained comparable to those in their control counterparts, suggesting that the shAMPKα1 cell clones maintain a relatively high capacity for AMPK signaling. **(c, d)** The impact of AMPKα1 downregulation on the response of RD cells to phenformin in terms of AMPK activation was further assessed by comparing the phenformin-induced increase in p-AMPKα1,2(T172) in shCTRL and shAMPKα1 cell clones. Normalized protein levels relative to the untreated controls detected in individual clones (c) or generally in the shCTRL and shAMPKα1 clone groups (d) are plotted as the mean ± SD. Statistical significance was determined by an unpaired two-tailed Student’s t-test (d), ns – not significant.

**Fig. S19:**
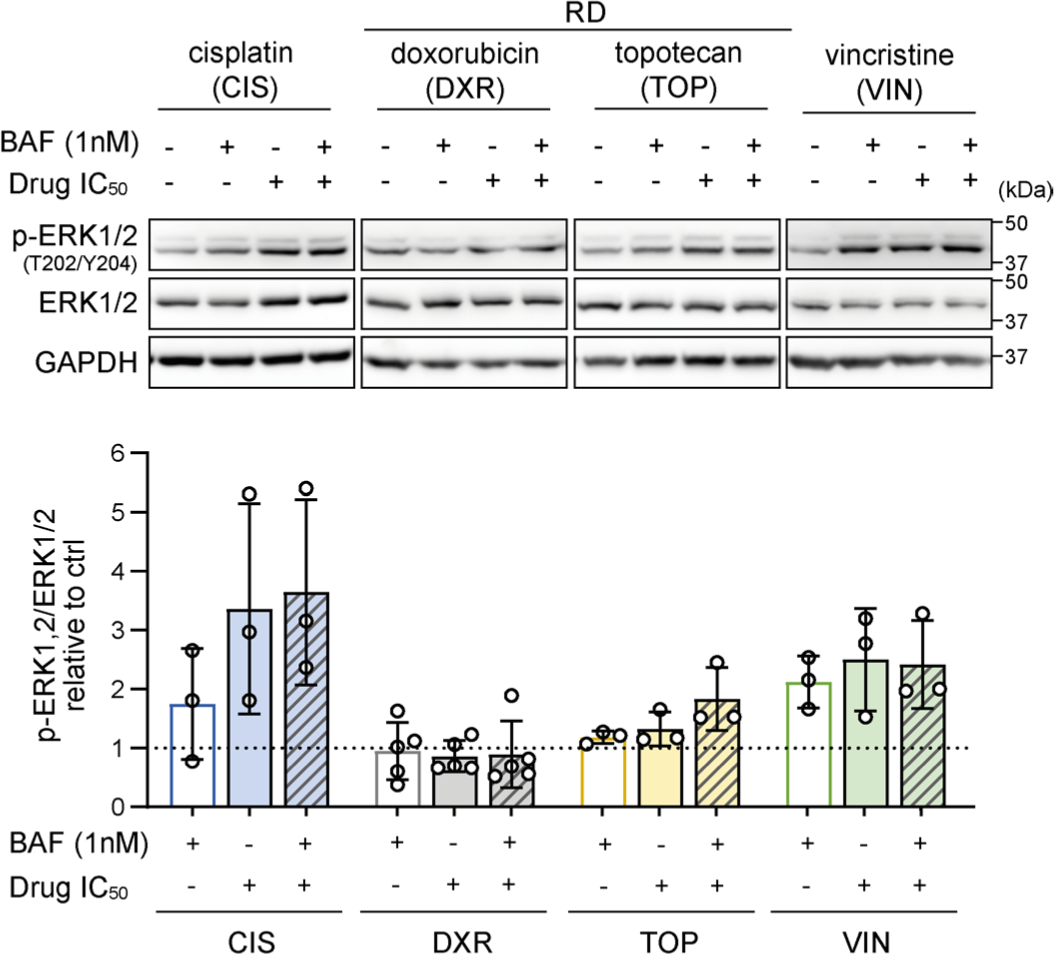
Chemotherapy-induced ERK activation does not correspond with chemotherapy-induced DRP1 activation. Western blotting detection and densitometric analysis of ERK kinase and its activated form, p-ERK1/2(T202/YT204), in RD rhabdomyosarcoma cells after 72-hour exposure to 1 nM bafilomycin A1 (BAF), chemotherapy drugs at the IC50 dose, or combinations of BAF and chemotherapy drugs revealed that cisplatin (CIS) and vincristine (VIN), but not doxorubicin (DXR) or topotecan (TOP), induce ERK activation. In contrast, all these chemotherapy drugs induced an increase in DRP1 activating phosphorylation (Fig. 2c, d). Normalized protein levels are plotted relative to the untreated controls as the mean ± SD. Statistical significance was determined by one-way ANOVA followed by Tukey’s multiple comparisons test.

**Fig. S20:**
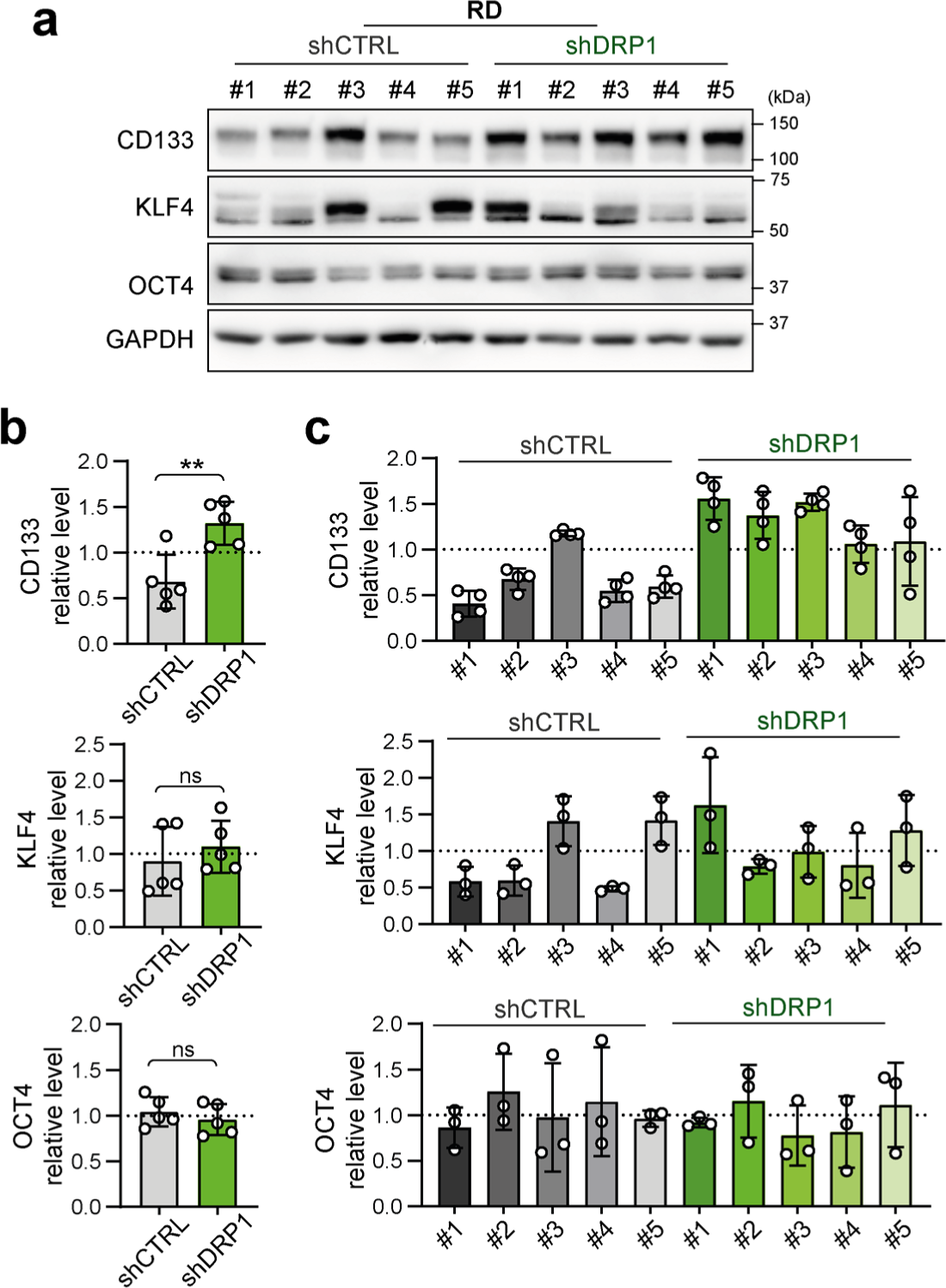
Stable knockdown of the mitochondrial fission mediator DRP1 did not decrease the expression of stemness-associated markers in RD rhabdomyosarcoma cells. **(a-c)** Western blotting (a) and densitometric analysis (b, c) of stemness-associated markers in individual RD rhabdomyosarcoma control (shCTRL) and DRP1-downregulated (shDRP1) cell clones. **(b-c**) Mean normalized protein levels detected in all shCTRL and shDRP1 clones (b) and biological replicates of individual clones (c) are plotted as the mean ± SD. Statistical significance was determined by unpaired two-tailed Student’s t-test, ns – not significant.

## SUPPLEMENTARY TABLES

**Supplementary Table 1.**
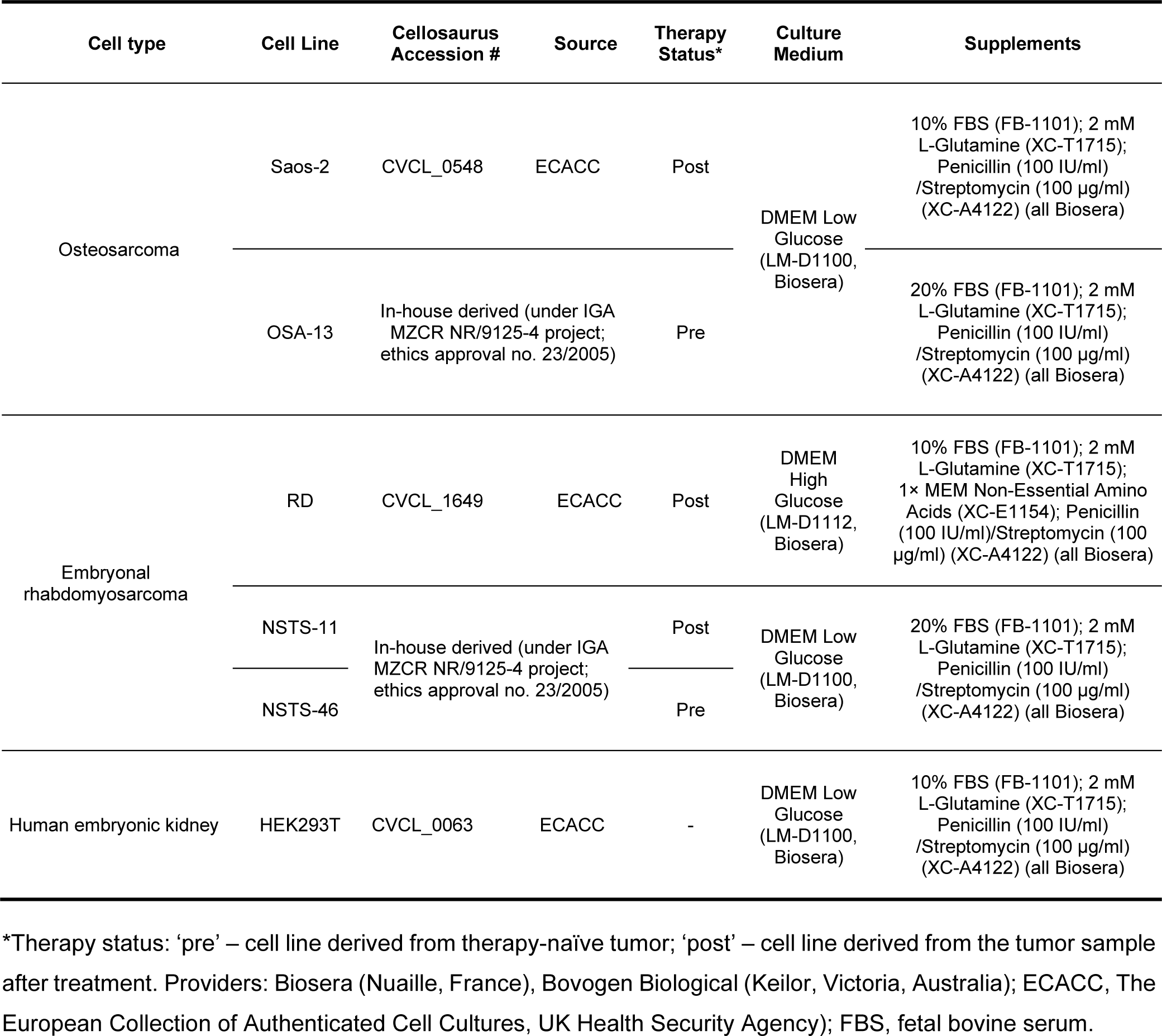
Summary of cell lines and culture media composition.

| Cell type | Cell Line | Cellosaurus Accession # | Source | Therapy Status* | Culture Medium | Supplements |
| --- | --- | --- | --- | --- | --- | --- |
| Osteosarcoma | Saos-2 | CVCL_0548 | ECACC | Post | DMEM Low Glucose (LM-D1100, Biosera) | 10% FBS (FB-1101); 2 mM L-Glutamine (XC-T1715); Penicillin (100 IU/ml) /Streptomycin (100 µg/ml) (XC-A4122) (all Biosera) |
|  | OSA-13 | In-house derived (under IGA MZCR NR/9125-4 project; ethics approval no. 23/2005) |  | Pre |  | 20% FBS (FB-1101); 2 mM L-Glutamine (XC-T1715); Penicillin (100 IU/ml) /Streptomycin (100 µg/ml) (XC-A4122) (all Biosera) |
| Embryonal rhabdomyosarcoma | RD | CVCL_1649 | ECACC | Post | DMEM High Glucose (LM-D1112, Biosera) | 10% FBS (FB-1101); 2 mM L-Glutamine (XC-T1715); 1× MEM Non-Essential Amino Acids (XC-E1154); Penicillin (100 IU/ml)/Streptomycin (100 µg/ml) (XC-A4122) (all Biosera) |
|  | NSTS-11 | In-house derived (under IGA MZCR NR/9125-4 project; ethics approval no. 23/2005) |  | Post | DMEM Low Glucose (LM-D1100, Biosera) | 20% FBS (FB-1101); 2 mM L-Glutamine (XC-T1715); Penicillin (100 IU/ml) /Streptomycin (100 µg/ml) (XC-A4122) (all Biosera) |
|  | NSTS-46 |  |  | Pre |  |  |
| Human embryonic kidney | HEK293T | CVCL_0063 | ECACC | - | DMEM Low Glucose (LM-D1100, Biosera) | 10% FBS (FB-1101); 2 mM L-Glutamine (XC-T1715); Penicillin (100 IU/ml) /Streptomycin (100 µg/ml) (XC-A4122) (all Biosera) |

\*Therapy status: ‘pre’ – cell line derived from therapy-naïve tumor; ‘post’ – cell line derived from the tumor sample after treatment. Providers: Biosera (Nuaille, France), Bovogen Biological (Keilor, Victoria, Australia); ECACC, The European Collection of Authenticated Cell Cultures, UK Health Security Agency); FBS, fetal bovine serum.

**Supplementary Table 2.** Drugs used in the study.

| Drug | Manufacturer | Catalog number | Solvent |
| --- | --- | --- | --- |
| ABT-737 | Abcam | ab141336 | DMSO |
| Bafilomycin-A1 | Sigma–Aldrich | 19-148 | DMSO |
| Cisplatin | Sigma–Aldrich | 232120 | PBS |
| Chloroquine | Tocris | 4109 | H<sub>2</sub>O |
| Doxorubicin | Tocris | 2252 | DMSO |
| Etoposide | Tocris | 1226 | DMSO |
| Phenformin hydrochloride | Sigma–Aldrich | P7045 | H<sub>2</sub>O |
| Sunitinib | Cell Signaling | 12328S | DMSO |
| Trametinib | APEXBIO | GEN1590424 | DMSO |
| Topotecan | Accord Healthcare | 44/322/12-C | H<sub>2</sub>O |
| Vincristine | Tocris | 1257 | DMSO |

Providers: Abcam (Cambridge, UK), Accord Healthcare (London, UK), APEXBIO (Houston, TX, USA), CST – Cell Signaling Technology (Danvers, MA, USA), EMD Millipore (Billerica, MA, USA), Sigma–Aldrich (St. Louis, MO, USA), Serva (Heidelberg, Germany), and Tocris (Bristol, UK).

**Supplementary Table 3.**
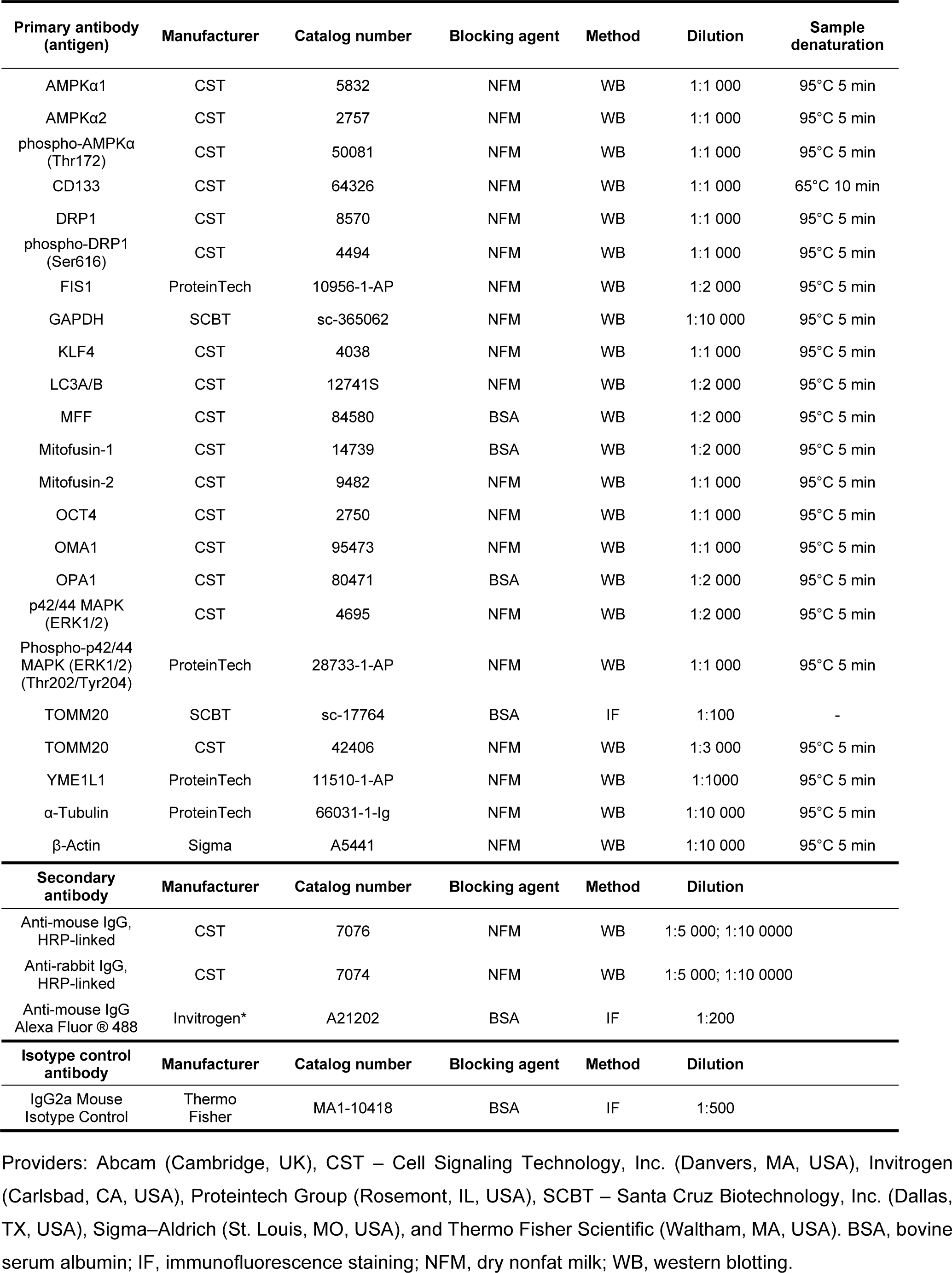
Antibodies used for immunodetection.

| Primary antibody (antigen) | Manufacturer | Catalog number | Blocking agent | Method | Dilution | Sample denaturation |
| --- | --- | --- | --- | --- | --- | --- |
| AMPKα1 | CST | 5832 | NFM | WB | 1:1 000 | 95°C 5 min |
| AMPKα2 | CST | 2757 | NFM | WB | 1:1 000 | 95°C 5 min |
| phospho-AMPKα (Thr172) | CST | 50081 | NFM | WB | 1:1 000 | 95°C 5 min |
| CD133 | CST | 64326 | NFM | WB | 1:1 000 | 65°C 10 min |
| DRP1 | CST | 8570 | NFM | WB | 1:1 000 | 95°C 5 min |
| phospho-DRP1 (Ser616) | CST | 4494 | NFM | WB | 1:1 000 | 95°C 5 min |
| FIS1 | ProteinTech | 10956-1-AP | NFM | WB | 1:2 000 | 95°C 5 min |
| GAPDH | SCBT | sc-365062 | NFM | WB | 1:10 000 | 95°C 5 min |
| KLF4 | CST | 4038 | NFM | WB | 1:1 000 | 95°C 5 min |
| LC3A/B | CST | 12741S | NFM | WB | 1:2 000 | 95°C 5 min |
| MFF | CST | 84580 | BSA | WB | 1:2 000 | 95°C 5 min |
| Mitofusin-1 | CST | 14739 | BSA | WB | 1:2 000 | 95°C 5 min |
| Mitofusin-2 | CST | 9482 | NFM | WB | 1:1 000 | 95°C 5 min |
| OCT4 | CST | 2750 | NFM | WB | 1:1 000 | 95°C 5 min |
| OMA1 | CST | 95473 | NFM | WB | 1:1 000 | 95°C 5 min |
| OPA1 | CST | 80471 | BSA | WB | 1:2 000 | 95°C 5 min |
| p42/44 MAPK (ERK1/2) | CST | 4695 | NFM | WB | 1:2 000 | 95°C 5 min |
| Phospho-p42/44 MAPK (ERK1/2) (Thr202/Tyr204) | ProteinTech | 28733-1-AP | NFM | WB | 1:1 000 | 95°C 5 min |
| TOMM20 | SCBT | sc-17764 | BSA | IF | 1:100 | - |
| TOMM20 | CST | 42406 | NFM | WB | 1:3 000 | 95°C 5 min |
| YME1L1 | ProteinTech | 11510-1-AP | NFM | WB | 1:1000 | 95°C 5 min |
| α-Tubulin | ProteinTech | 66031-1-Ig | NFM | WB | 1:10 000 | 95°C 5 min |
| β-Actin | Sigma | A5441 | NFM | WB | 1:10 000 | 95°C 5 min |
| Secondary antibody | Manufacturer | Catalog number | Blocking agent | Method | Dilution |  |
| Anti-mouse IgG, HRP-linked | CST | 7076 | NFM | WB | 1:5 000; 1:10 0000 |  |
| Anti-rabbit IgG, HRP-linked | CST | 7074 | NFM | WB | 1:5 000; 1:10 0000 |  |
| Anti-mouse IgG Alexa Fluor® 488 | Invitrogen* | A21202 | BSA | IF | 1:200 |  |
| Isotype control antibody | Manufacturer | Catalog number | Blocking agent | Method | Dilution |  |
| IgG2a Mouse Isotype Control | Thermo Fisher | MA1-10418 | BSA | IF | 1:500 |  |

Providers: Abcam (Cambridge, UK), CST – Cell Signaling Technology, Inc. (Danvers, MA, USA), Invitrogen (Carlsbad, CA, USA), Proteintech Group (Rosemont, IL, USA), SCBT – Santa Cruz Biotechnology, Inc. (Dallas, TX, USA), Sigma–Aldrich (St. Louis, MO, USA), and Thermo Fisher Scientific (Waltham, MA, USA). BSA, bovine serum albumin; IF, immunofluorescence staining; NFM, dry nonfat milk; WB, western blotting.

